# Inoculum-to-interface distance tunes productivity in static liquid cultures of wood-decay fungi

**DOI:** 10.64898/2026.10.13.758717

**Authors:** Piet Dimitriadis, Anouk Verstuyft, Jara Saluena Martin, Rafael Philippini, Annah-Ololade Sangosanya, Simon Vandelook, Wim De Malsche, Eveline Peeters

## Abstract

Static liquid cultures of wood-decay fungi underpin applications in materials, food, and medicine, yet the mechanisms governing pellicle formation remain poorly understood, limiting rational optimisation at scale. We developed a time-lapse scanning and image analysis platform to visualise the process of pellicle formation across five white-rot fungal species, of which *Trametes versicolor* formed the thickest, heaviest pellicles and completed formation fastest. Kymographs revealed a reproducible four-stage sequence after inoculation and sedimentation of the biomass: recovery of viability, upward growth, pellicle initiation, and thickening. We formalised this as a model in which the time to pellicle establishment is determined by the distance between the settled biomass and the liquid–air interface. Increasing liquid depth relative to a fixed inoculum loading raised biomass and mat thickness, but prolonged formation. The results identify surface proximity as the dominant predictor of initiation timing. By fixing the inoculum at the liquid-air interface with a membrane, we validated that minimising the distance between the sedimented mycelium and the liquid surface accelerated and stabilised formation without compromising on pellicle biomass. By exposing the most relevant control variables, this work provides a quantitative basis for designing reliable liquid-state surface fermentations for emerging applications involving mycelium.

## Introduction

Fungal mycelium, particularly from wood-decay Basidiomycota, underpins diverse biotechnological applications, from food fermentations and secondary metabolite production to emerging mycelium-based materials(1–3). Pellicles formed during liquid-state surface fermentation (LSSF) can be developed into renewable alternatives to leather, meat, and packaging, with further potential in wastewater remediation and biomedical scaffolding (4–6). Yet how fungi colonise a liquid surface and form into a pellicle remains mechanistically unclear, constraining systematic process optimisation and the scalability of this fermentation mode.

Wood-decay fungi are well suited to this route. Many form mechanically robust mycelium underpinned by differentiated hyphal architectures. The trimitic systems of Polyporales such as *Ganoderma lucidum* and *Trametes versicolor*, combine generative, skeletal, and binding hyphae to produce remarkably durable tissue (7). In practice, however, static cultivation of mycelium for materials production is held back by biological stochasticity (8), phenotypic diversity (9,10), and susceptibility to contamination (11).

Surface films are not unique to filamentous fungi: pellicles of bacterial cellulose, yeast, and *B. subtilis* also assemble at air–liquid interfaces through comparatively well-characterised pathways (12–14). However, no equivalent account exists for how a dispersed suspension of mycelium reorganises and grows into a coherent film. Although tropic responses such as aerotropism and chemotropism have been described in filamentous fungi (15,16), and hydrophobin self-assembly is known to lower water surface tension sufficiently for hyphae to breach the liquid surface (17), neither has been connected to how a dispersed mycelial suspension first reaches the interface in static liquid culture.

Here, we introduce a time-lapse scanning and image-analysis method to follow LSSF cultures of wood-decay fungi *in situ* and apply it across a five-species screen to resolve how pellicles form. Pellicle development follows a reproducible sequence: the inoculum sediments, the culture recovers, and hyphae extend upwards to the liquid–air interface. Through statistical analysis, we unveil that increasing distance to the interface increases variance in pellicle initiation times. By revealing this critical variable, this work provides a quantitative basis for tuning the rate and reproducibility of fungal static cultures both in the lab, and at scale.

## Results

### Timelapse image analysis reveals static culture stages

We investigated how fungal pellicles arise from dispersed mycelium in static liquid culture. To characterise the formation kinetics, we developed a time-resolved image analysis method that converts sequential images into kymographs, recording how the colony front moves through space over time (**Figure 1A**). Kymographs constructed from scans of 2 mL cultures containing *T. versicolor* (**Figure 1B**) reveal a consistent formation pathway illustrated in **Figure 1C**. Following inoculation, mycelial fragments sedimented to the vessel bottom. After a recovery period, a coherent growth front developed and advanced upward through the liquid column. Pellicles formed when the upward-growing hyphae reached the liquid–air interface and subsequently thickened in the oxygen rich environment.

**Figure 1.**
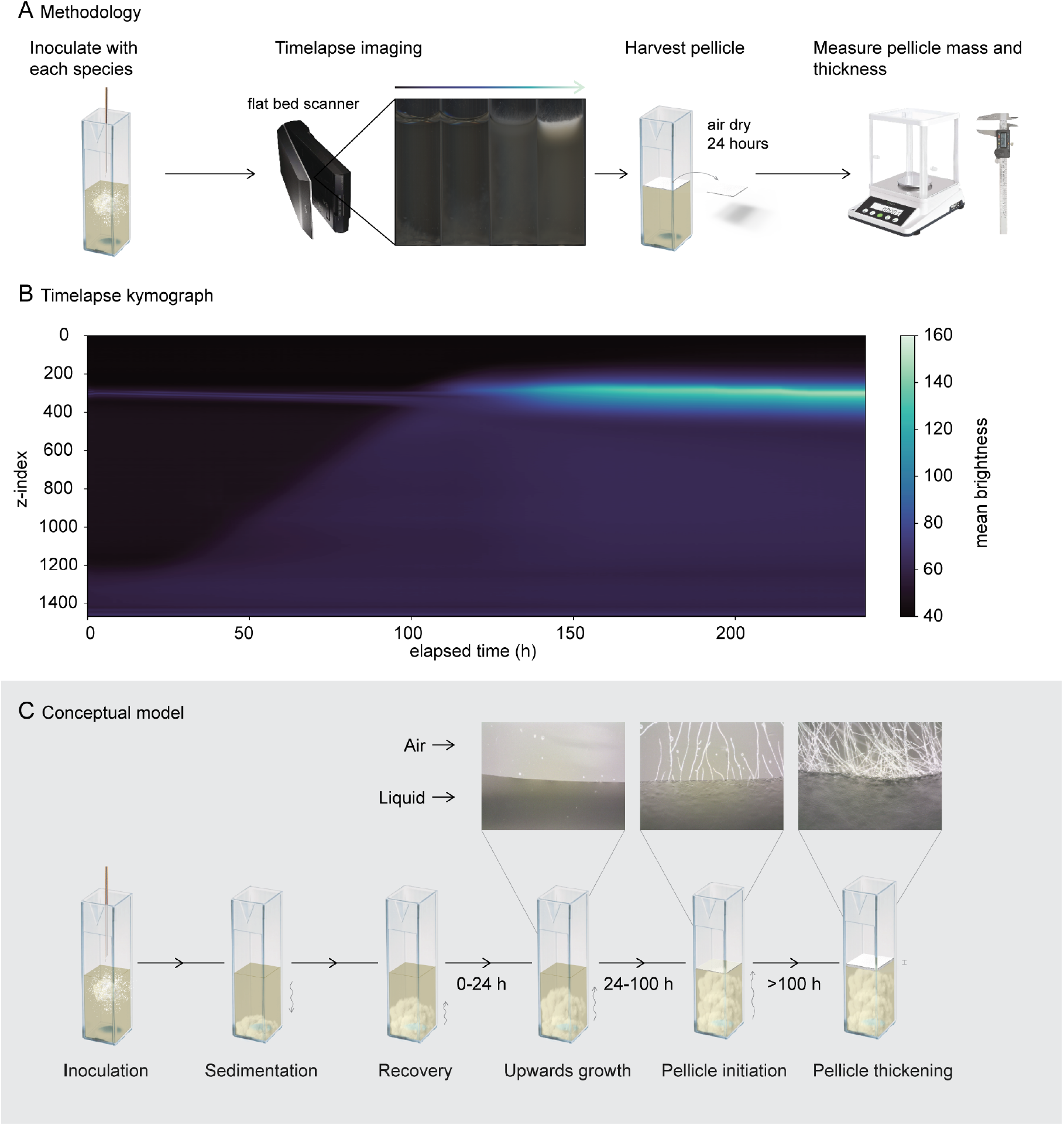
Timelapse image analysis reveals four stages of static culture progression. **A**) Methodology used for quantifying growth kinetics and pellicle properties. **B**) Kymographs reveal spatial distribution of mycelium in cuvettes through time. Brighter regions indicate the presence of mycelium. Narrow band at z≅ 300 px is the liquid-air interface which brightens as the white pellicle forms and thickens **C**) Conceptual model of static liquid culture progression. Following inoculation, mycelium sediments to the vessel base, and undergoes a short recovery phase. It is followed by a growth front then extends upwards through the liquid column until it reaches the liquid-air interface, where pellicle formation begins; the pellicle subsequently thickens into a cohesive mat.

Kymographs generated distinctly brighter pixels in regions where mycelium was present. By averaging the pixels in each row of a cuvette, the vertical position of the extending mycelium colony was traceable through time. The lag phase duration, here defined as the time it takes for the pellicle to initialise, was determined by averaging the rows of pixels around the liquid-air interface and fitting a modified Gompertz model with the lag phase parametrised (18). The time to stationary phase, here defined as the point at which the pellicle stops growing, was calculated by taking the time corresponding to 95% of the model amplitude.

We constructed the lag phase duration as the sum of recovery and upwards growth durations. Since sedimentation occurred prior to the first scan, and in a time frame smaller than what was resolvable with the scanner setup, it was not included with the lag phase durations determined through image analysis.

### Growth dynamics vary by species

Five phylogenetically diverse, mushroom forming wood-decay fungi were screened: *Trametes versicolor (T. versicolor), Ganoderma sessile (G. sessile), Fomes fomentarius (F. fomentarius), Pleurotus ostreatus (P. ostreatus),* and *Schizophyllum commune (S. commune)* (**Figure 2**). Kymographs for each species (**Supplementary Figures S1-S20**) depict the culture progression across 10 days. Post-hoc image analysis enabled quantification of culture kinetics, while measuring thickness and mass were performed on the air-dried pellicles (**Figure 2A-D**).

**Figure 2.**
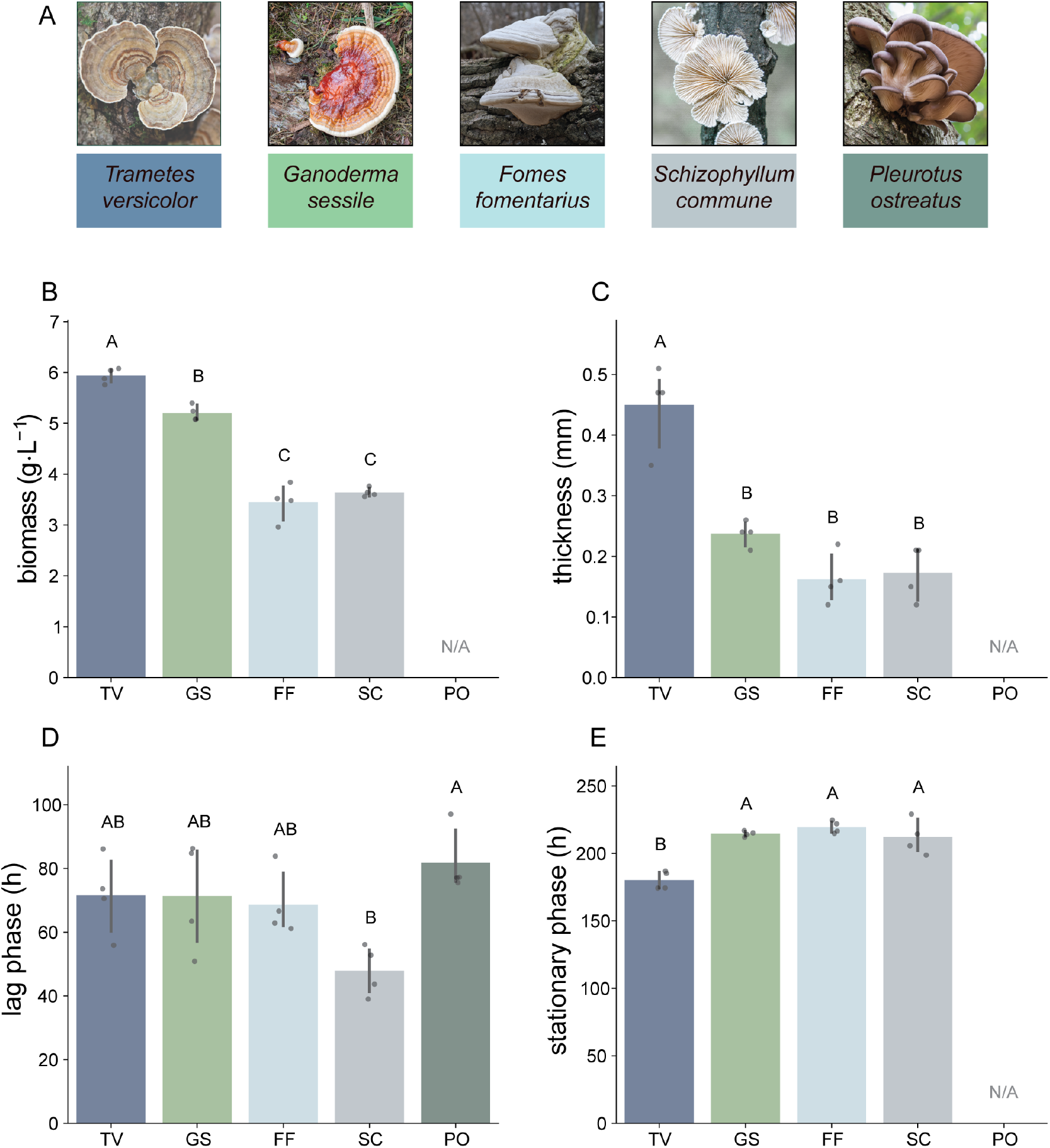
Species selection determines fermentation performance. **A**) Five wood decay fungi were studied via timelapse scanning in cuvettes; **B**) pellicle yields; **C**) pellicle thicknesses; **D**) lag-phase durations; and **E**) time to stationary phase (cessation of pellicle thickening) for each species. Shared letters above bars indicate species that do not differ significantly: biomass, pellicle thickness, and lag-phase duration were compared by one-way ANOVA with Tukey HSD post hoc tests, and time to stationary phase by Welch’s ANOVA with Games–Howell post hoc tests (two-sided, α = 0.05, n=4). Images of fungi were sourced from Adobe Stock.

Biomass yield differed strongly across species (**Figure 2A**) (one-way ANOVA, F(3, 12) = 126.31, *p* < 0.05, partial η² = 0.97). Tukey HSD post hoc comparisons (*n* = 4 per group) showed that *T. versicolor* produced significantly greater biomass yield than *S. commune* (*p* < 0.05 g = 16.5), *F. fomentarius* (*p* < 0.05, g = 7.79), and *G. sessile* (p < 0.05, g = 4.27). *G. sessile* also exceeded both *S. commune* (*p* < 0.05, g = 10.91) and *F. fomentarius* (*p* < 0.05, g = 5.44), while no significant difference was detected between *S. commune* and *F. fomentarius* (*p > 0.05*). Pellicles formed by *P. ostreatus* could not be handled reproducibly and were excluded from biomass analysis.

Pellicle thickness followed a similar pattern (**Figure 2B**) (one-way ANOVA, F(3, 12) = 31.79, *p < 0.05*, partial η² = 0.89). Tukey HSD post hoc comparisons (*n* = 4 per group) showed that *T. versicolor* produced significantly thicker pellicles than *F. fomentarius* (*p* < 0.05, g = 4.37), *S. commune* (*p* < 0.05, g = 4.13), and *G. sessile* (*p* < 0.05, g = 3.62), whereas differences among the latter three species were not statistically significant (all *p* > 0.05).

*S. commune* exhibited a shorter lag phase than *P. ostreatus* (p < 0.05, g = 3.22, *n* = 4 per group), representing a very large effect size. Differences between *S. commune* and the remaining species were not statistically significant (*p* > 0.05) (**Figure 2C**), although several comparisons showed large effect magnitudes (Hedge’s g ≈ 1.52–1.97). This behaviour coincided with spontaneous upward displacement of the hyphae (**Supplementary Figures S13-S16**), though it may alternatively reflect a higher intrinsic tip extension rate.

Time to stationary phase differed across species (**Figure 2D**) (Welch’s ANOVA, F(3, 5.77) = 28.17, *p* < 0.05, partial η² = 0.84). Games–Howell post hoc comparisons (*n* = 4 per group) showed that *T. versicolor* reached stationary phase significantly earlier than *F. fomentarius* (*p* < 0.05, g = 5.90), *G. sessile* (*p* < 0.05, g = 5.96) and *S. commune* (*p* < 0.05, g = 2.65). *P. ostreatus* was excluded from this analysis as cultures never reached stationary phase during the studied timeframe.

Following inoculation, hyphae from all species initially appeared to sediment and subsequently grew upward toward the liquid surface. Some species however, showed this feature less distinctly, with multiple apparent initiation points visible throughout the vertical axis of the liquid column after a recovery period (**Supplementary Figures S5-S12**). *S. commune* showcased a distinct behaviour in which previously submerged mycelium rapidly displaced upward toward the liquid–air interface after growing for approximately 80 hours (**Supplementary Figures S13-S16**).

Recovery time was scored manually from each cuvette’s kymograph as the elapsed time at which the sedimented mycelial front began sustained upward extension (*n* = 4 cuvettes per species). Group means are shown with 95 % confidence intervals (**Supplementary Figure S21**). A Brown–Forsythe test confirmed unequal variances across species (F = 6.80, *p* < 0.05), so species were compared with Welch’s ANOVA. Recovery time differed significantly among species (Welch F(4, 6.90) = 14.05, *p* < 0.05, η² = 0.55), with *P. ostreatus* recovering earliest (20.4 ± 1.0 h) and *G. sessile* latest (35.5 ± 5.0 h). The surviving pairwise contrasts were *G. sessile* versus *P. o*streatus (*p* < 0.05) and *T. versicolor* versus *P. ostreatus* (*p* < 0.05).

Overall, *T. versicolor* emerged as the most performant pellicle-producing strain under the tested conditions, combining rapid progression to stationary phase with the highest biomass yield and pellicle thickness, while progressing to the stationary phase most consistently.

### Interface proximity governs rate of production

We investigated the effect of incremental liquid levels on pellicle formation dynamics and biomass productivity in cultures of *T. versicolor*. Pellicles emerged sooner in shallower cultures (**Figure 3A**) where the distance between the sedimented mycelium and the liquid air interface (Δ*z*) was smaller, visible in the culture kymographs (**Supplementary Figures S22-S41**). Lag phase duration and time to stationary phase both increased linearly with Δ*z* (r^2^ = 0.905, *n* = 20 and r^2^ = 0.902, *n* = 20 respectively), while biomass and thickness followed saturating trends (**Figure 3B**). These results indicate that distance between the sedimented mycelium and liquid air interface is a dominant predictor of pellicle initiation timing.

**Figure 3.**
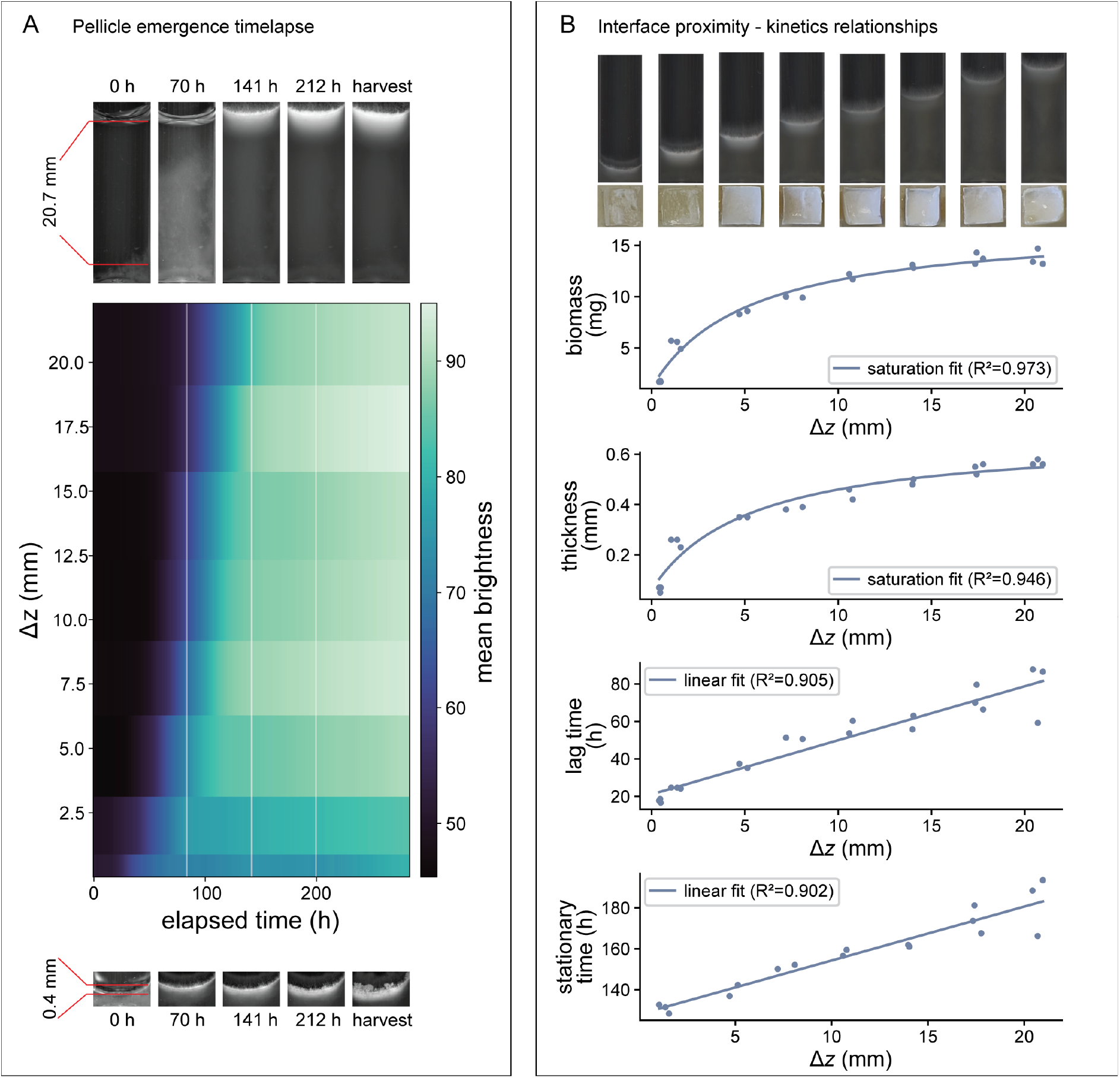
Liquid depth establishes a trade-off between rate of production and pellicle yield. Cuvettes of increasing liquid volume relative to a fixed inoculum volume were scanned over 10 days. **A**) Pellicle emergence time increased with Δz. Brighter regions indicate the presence of mycelium at the liquid air interface. **B**) Pellicle mass reached saturation at higher liquid levels, while lag and stationary time grew linearly (n = 2–3 independent cultures per depth).

Variability between replicates also increased with liquid level. The between-replicate standard deviation in lag time rose from ≈1 h at the shallowest fill (1.8 mm) to ≈16 h at the deepest (21.4 mm) (Spearman ρ = 0.88, *p* <0.005, n = 8 levels), and the coefficient of variation rose in parallel (ρ = 0.79, p < 0.05). Classical heteroscedasticity tests on the individual cultures were not significant (Breusch–Pagan *p* > 0.05; Goldfeld–Quandt *p* > 0.05), consistent with low power at *n* = 20; we therefore fitted a Bayesian model separating measurement uncertainty from biological variability, which assigned posterior probability 0.998 that biological variability in lag time increases with Δ*z* (γ₁ = 0.67, 94% HDI [0.18, 1.17]). Deeper liquid columns therefore delayed pellicle initiation and made it markedly less reproducible.

Recovery times reduced with increasing Δ*z* (*p* < 0.05), although no clear trend was determined. Taken together, increasing liquid depth systematically shifts pellicle biomass and thickness higher, but at the cost of slower and less reliable kinetics, establishing a trade-off between pellicle productivity and process reliability.

### Membranes accelerate pellicle formation without a<ecting biomass

Introducing a membrane scaffold to promote early interfacial proximity of submerged *T. versicolor* mycelium (**Figure 4A**) accelerated fermentation kinetics visible in the culture kymographs and growth curves (**Supplementary Figures S43-63**). Cultures grown with membranes exhibited substantially shorter lag phases and reached stationary phase earlier than controls (**Figure 4C** and **D**) without compromising the overall productivity (**Figure 4E**).

**Figure 4.**
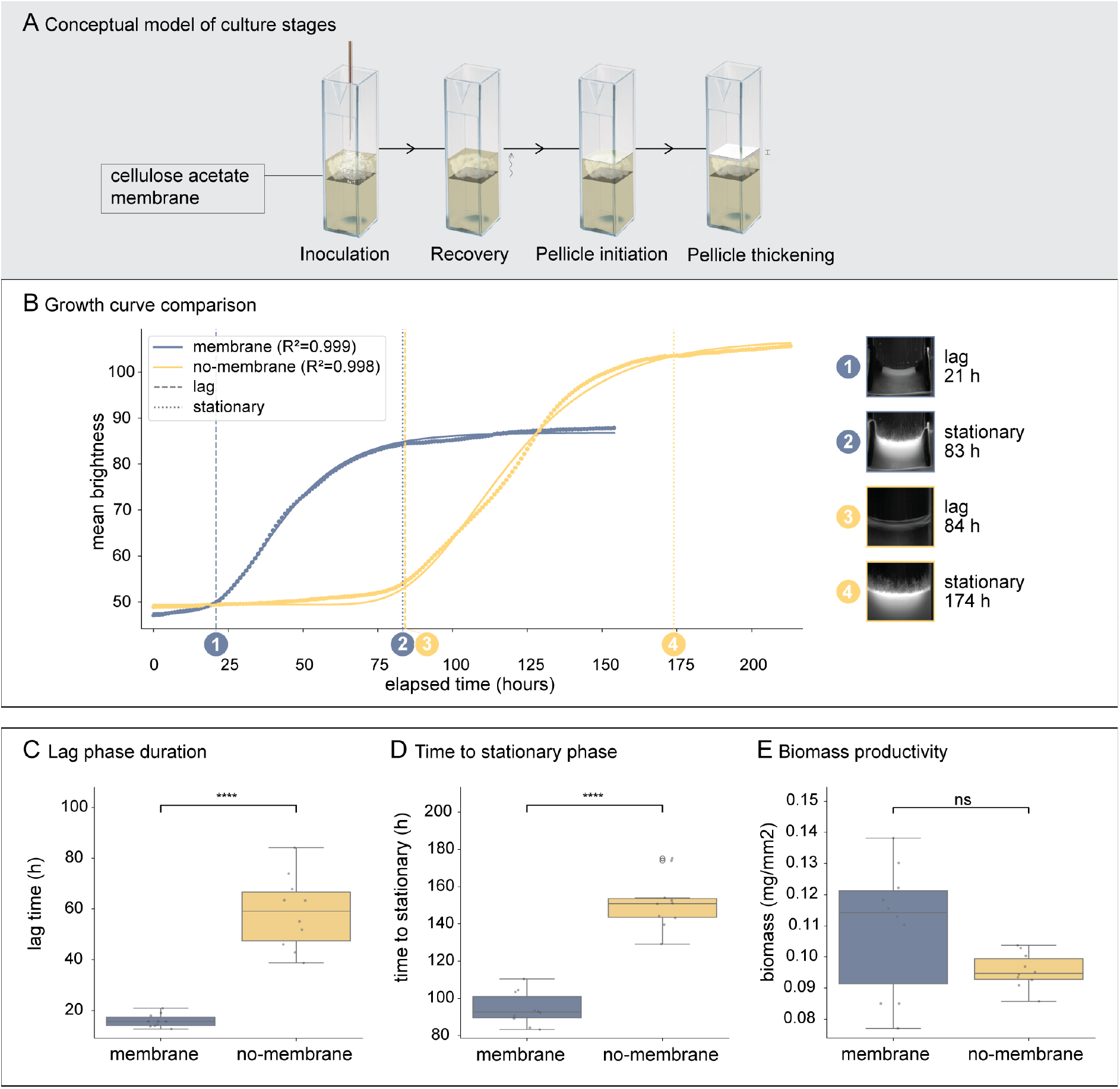
Membranes decrease the distance between sedimented inoculum and the interface, accelerating pellicle formation. **A**) Conceptual model of culture progression in a membrane-supported culture: a cellulose-acetate membrane holds the inoculum at the liquid-air interface, removing the upward growth stage required in control cultures, reducing the time to pellicle initiation**. B**) Representative growth curves for a membrane-supported culture and an unsupported, no-membrane control. Dots measured intensity; solid line, fitted Gompertz model. Lag-phase duration and time to stationary phase were extracted from these fits. Absolute brightnesses do not correlate with biomass yields. **C**) Distribution of lag-phase durations for cultures with and without a membrane **D**) Distribution of time taken to reach stationary phase for cultures with and without a membrane. **E**) Distribution of dry biomass per surface area for cultures with and without a membrane). Box plots show the median (centre line), interquartile range (box), and 1.5× IQR (whiskers); points are individual cultures.

Lag-phase duration was significantly reduced by 42.75 h (Welch’s ANOVA, *p* < 0.005) with a concurrent 5.6-fold reduction in standard deviation (Brown–Forsythe *p* < 0.005; Fligner– Killeen *p* = < 0.005). Time to stationary phase also significantly shortened by 56.93 h (Welch’s ANOVA, *p* = < 0.005), although the associated reduction in standard deviation (1.60×) was not statistically significant (Brown–Forsythe *p* > 0.05). Despite these kinetic improvements, no significant difference in biomass productivity was observed between conditions (*p* > 0.05), indicating that the membrane primarily affects the rate, rather than the final biomass accumulation.

Biomass variance increased in membrane-supported cultures (Brown–Forsythe *p* = 0.05; Fligner–Killeen *p* < 0.05). This likely reflected the additional colonisable surface area introduced by the textured 3D-printed membrane scaffold. Furthermore, trapped air bubbles beneath the membrane were observed to expand the available growth interface (**Supplementary Figure S42**). For productivity measurements (**Figure 4E**), only pellicle-associated biomass was quantified; however, distinguishing pellicle from non-pellicle biomass was not always unambiguous, introducing a potential source of error. Nevertheless, the principle remains clear, earlier surface contact promotes faster pellicle initiation.

## Discussion

### Pellicle initiation is growth driven

The live imaging methodology used in this study revealed how submerged mycelium extends to the liquid surface. The continuous, front-like advance of biomass in the vertical column indicates that pellicle initiation is consistent with directed growth, rather than transport mechanisms other interfacial colonisers rely on. Motile bacteria such as *B. subtilis* swim to the interface by flagellar motility and aerotaxis (13); yeasts rise through CO_2_-driven buoyancy and flocculation (14); and bacterial-cellulose producers, being obligate aerobes, are carried to and held at the oxygen-rich interface by the crystalline cellulose they secrete (19). In each of these, the cells are carried to the surface. Filamentous mycelium cannot be carried since the fragments are non-motile, denser than the medium (20), and do not trap CO_2_ like with yeast, thus the only route to the interface is to grow across the intervening distance. This is what makes pellicle initiation in filamentous fungi a fundamentally growth-limited process and, as we experimentally validated, one whose timing is governed by the distance between the sedimented inoculum and the liquid surface Δ*z*.

### Interface distance determines both the timing and reproducibility of pellicle initiation

As liquid level increases, the law of large numbers would suggest that variances in moment-to-moment colony extension rate should average out, and less variance in pellicle initiation time would be observed. However, colony growth rate is a function of the species and the medium, but also of local changes in gene expression. These deviations arise from two distinct sources: heterogeneity in the biological state of the inoculum fragments, and the stochasticity of tip growth itself.

It is likely that growth rate is tied to the biological state of the mycelium in the inoculum, related to its age and location in the preculture colony. According to a study by Wösten et al., approximately 50% of variation in gene expression comes from the medium, and the other 50% can be attributed to differentiation in the vegetative mycelium (21). In practice, this is why mycelium is usually taken from the edge of an agar culture rather than the centre when subculturing: inocula from the periphery appear to grow faster and give more reproducible results. This idea is cogent based on our understanding of cytoplasm streaming toward the growing tip, which leaves hyphae behind the peripheral growth zone progressively vacuolated and depleted of mobile cytoplasm (22). Excised tissue that is cytoplasm-depleted shows a measurable lag before resuming net extension, tied to the time needed to mobilise internal reserves (23). Although whether the degree of depletion in the excised piece scales with the length of that recovery phase has not been directly tested to our knowledge.

Inoculum produced from blending a whole colony, or several, contains a mixture of these biological states. When aliquots are transferred into their containers, the distribution of these states is not necessarily consistent between cultures or evenly distributed within a culture. This provides a probable source for the growth rate variances observed in this study.

However, differences in cytoplasm availability in early cultures is not the only factor which might contribute to growth rate variance. Genes relating to branching, cell wall remodelling, and enzyme production might also be critical. Branching governs the size of the tip population, and therefore the number of concurrent trajectories exploring the liquid. As such, the time to interface colonisation also depends on the relative contribution of either increasing search parallelism (branching) or single tip extension efficiency (non-branching). While increased branching increases the number of exploratory tips, it may also alter extension dynamics through resource allocation and structural constraints. This effect depends on branching mode. In *Neurospora crassa*, lateral branching introduces new tips without necessarily interrupting extension of the parent hypha, whereas apical branching redistributes growth at the leading edge and can reduce forward extension of individual tips (24).

The growth dynamics of colonies are also governed by biochemical signalling and geometric constraints. Positive aerotropism biases growth towards the liquid air interface (16), while directional persistence limits reorientation (25), producing correlated trajectories. Local interactions further constrain growth: negative autotropism diverts tips away from neighbouring hyphae (15), and impermeable boundaries (e.g., vessel walls) restrict available trajectories and biases growth away from obstructed regions (15,26).

Combined, the discussed effects define a stochastic process that is highly sensitive to culture geometry, the biological state of the inoculum fragments, and their spatial distribution. As Δ*z* increases, the longer growth period and greater spatial domain allow local deviations in growth rate to compound and accumulate, thereby increasing variability in the measured pellicle initiation timing. Further investigation should establish the relative contributions of these factors in static cultures of wood-decay fungi. Research analysing gene expression associated with differentiation and tissue-specific growth rates could be particularly illuminating.

### Liquid height acts on yield and growth rate in opposite directions but the two are separable

Increasing liquid level elevates the nutrients available to a growing pellicle, enabling higher biomass accumulation up until oxygen becomes limiting. This phenomenon might also explain why recovery times were shorter in higher liquid levels compared to the lowest liquid level, where more nutrients were available for cell repair. However, increasing liquid level for a fixed inoculum ratio extends lag phase duration, risking contamination in the early stages of a culture.

There are a few solutions to this trade-off such as increasing inoculum loading, raising soluble nutrient concentration, or opting for an insoluble carbon source and loading it in excess. Increasing the inoculum loading increases the height of the sedimented mycelium and lowers the distance between it and the liquid surface. Since the biomass in the inoculum occupies very little volume, this does not greatly lower the available nutrients.

Increasing nutrient concentration while simultaneously lowering the liquid level enables a reduction in the distance between the inoculum and liquid surface without reducing the total moles of nutrients available for fermentation. Thus, the pellicle biomass should not change significantly and might even increase as the fermentation becomes more efficient. However, increasing the inoculum loading may become uneconomical at scale by necessitating greater upstream fermentation costs. Additionally, increasing nutrient concentration to lower the liquid level may introduce osmotic stress, altering hyphal cell wall regulation (27) and subsequent material properties.

Utilising an insoluble carbon source, paste and slurry fermentation techniques (28) have resolved this trade-off, and feature more sustainable upstream processing. In this fermentation mode the inoculum can be positioned at the slurry-air interface where its sedimentation is impeded by the packed bed of substrate particles.

A similar effect was observed in this study where fermentation time was reduced when using a membrane scaffold which increased the proximity of the inoculum to the liquid surface. By positioning biomass closer to the liquid–air interface, the membrane effectively decouples growth kinetics from pellicle yield, enabling both rapid colonisation and pellicle mass to reach saturation. As discussed earlier, in the absence of a membrane, growth kinetics are inherently stochastic, terminating only when a hyphal filament reaches the interface and initiates pellicle formation. Constraining the vertical distance between sedimented biomass and the interface reduces the spatial domain of the culture, thereby limiting deviations in initiation timing. By extension, strategies that promote early surface contact may also limit pellicle anisotropy in larger, sparsely inoculated cultures, although this was not demonstrated here.

### Species alter the timing and reliability of initiation

The conceptual model devised here was consistent across the five species studied, with only minor variations observed in their respective kymographs. Possible causes for these observations are out-of-focal-plane growth or adhesion of small hyphal fragments to the cuvette wall shortly after inoculation. The five species progressed through the stages at different rates and produced materials of varying quality.

Delayed pellicle formation observed in *P. ostreatus* was likely related to the selected environmental conditions. Some *P. ostreatus* variants are known to prefer lower temperatures for mycelium development and fruiting compared to the other species screened. Additionally, a recent review by Rovers and Wösten indicated no instances where *P. ostreatus* was used to produce pure mycelium materials via LSSF (29), highlighting that LSSF is likely incompatible with this species.

A plausible explanation for the abrupt restructuring of submerged *S. commune* mycelium involves the physicochemical properties of the exopolysaccharide schizophyllan, which the organism produces in large quantities during liquid culturing (30,31). Schizophyllan is a β-(1→3)(1→6)-glucan that adopts a triple-helical conformation in water and behaves as a semiflexible rod. These rod-like polymers can undergo lyotropic liquid-crystal transitions above a critical concentration, and purified schizophyllan solutions show birefringence and cholesteric ordering at high concentrations (32). If schizophyllan accumulated around the mycelium to a level approaching this threshold, the resulting change is structure could induce a phase separation, explaining the sudden upwards displacement. However the concentrations required for this transition in purified solutions may exceed those typical of culture broth, and the phase behaviour of schizophyllan in complex media is not established. As no direct phase characterisation was performed here, this mechanism remains speculative; alternatives such as gas entrapment reducing buoyancy cannot be excluded.

Filamentous fungi are well known to adopt species-dependent macromorphologies in submerged culture. This includes freely dispersed mycelium, loose clumps, and dense pellets, governed by both genetics and cultivation conditions (33,34) with direct consequences for productivity (35). Our screen shows that this species dependence extends to liquid surface culture: under identical conditions, the five wood-decay species spanned the full range from robust, reproducible pellicle formation to effectively none. *T. versicolor* lay at the favourable end, generating the highest biomass, thickest pellicles, and fastest progression to stationary phase, while maintaining reliable kinetics. Whereas *P. ostreatus* formed pellicles that could not be handled reproducibly and never reached stationary phase within the experiment duration. Rather than a fixed ranking of organisms, these differences are consistent with the known sensitivity of hyphal development to environmental conditions, combined with the ecophysiological diversity of species adapted to distinct ecological niches (34,36,37). Given the better performance of *T. versicolor* under the conditions tested here, it was used for the following experiments. However, the other organisms we studied may occupy distinct niches within mycelium materials research, especially if conducted under their favourable process conditions. As such, it is paramount that a wider screening of species be performed to capture these phenotypic differences and provide mechanistic insights linking phylogenetic, ecological and biochemical diversity to performance under defined process conditions.

## Conclusion

We developed a time-lapse scanning and image-analysis method and used it to provide the first systematic model, to our knowledge, describing the stages of pellicle formation in static liquid cultures of wood-decay fungi. Using this approach, we found that increasing liquid depth raised biomass yield but at the expense of prolonged fermentation and less reliable pellicle initiation. A membrane-separated strategy substantially mitigated this trade-off, lowering both duration and kinetic variation of the fermentation without compromising biomass yield. Modelling of the system revealed that the distance between the sedimented mycelium and the liquid–air interface is the dominant predictor of both fermentation kinetics and reproducibility in liquid surface cultures of Basidiomycetes, providing a quantitative account for the behaviours observed across conditions.

Together, these findings and the methodologies underpinning them may inform the rational design of sustainable fungal bioprocesses to produce materials, food, and metabolites. Future developments should aim to localise inoculum at or close to the liquid surface without compromising material quality, where membranes, increased biomass loadings, insoluble feedstocks, and novel bioreactor designs all represent promising avenues.

## Materials and Methods

### Culture medium

Complete yeast medium (CYM) contained 20 g L⁻¹ D-glucose monohydrate, 3 g L⁻¹ peptone, 2 g L⁻¹ yeast extract, and 0.5 g L⁻¹ MgSO₄·7H₂O (all Duchefa Biochemie, Haarlem, Netherlands). To avoid Maillard reactions and caramelisation of the carbon source during sterilisation, glucose was autoclaved separately from the other components: D-glucose was dissolved in half the final volume of deionised water, and peptone, yeast extract, and MgSO₄·7H₂O in the remaining half. Both fractions were autoclaved (121°C, 15 min), cooled to room temperature, and combined aseptically. The pH of the combined medium was 6.5 and was not adjusted. Solid CYM for plate culture was supplemented with 2% w/v agar (Duchefa Biochemie, Haarlem, Netherlands).

### Fungal strains and inoculum preparation

Five wood-decay basidiomycete species were used: *Trametes versicolor* VUB-SV1 (38), *Schizophyllum commune* H4-8 (FGSC no. 9210), *Fomes fomentarius* (Mycelia.be, M9740), *Ganoderma sessile (*Mycelia.be, M9762), and *Pleurotus ostreatus* (Mycelia.be M2191). Strains were maintained on solid CYM at 27 °C. For each species, a 1 cm-diameter plug was excised from the actively growing colony margin with a sterile cork borer and transferred to a 50 mL centrifuge tube containing 20 mL CYM. Tubes were incubated statically at 27°C for ∼5 d until pellicle formation.

Each pellicle was homogenised at 8,000 rpm for ∼30 s (Biobase BH-50P; Biobase, Jinan, China), transferred to a 500 mL Erlenmeyer flask, and brought to 200 mL with fresh CYM. Submerged cultures were grown at 30°C with orbital shaking (180 rpm, 25 mm orbit) for 10 d to generate pelleted biomass. Pellets were stored in sterile 50 mL centrifuge tubes at 4°C for up to 1 month. Before use, biomass was pelleted (9,500 x g, 5 min), resuspended in fresh CYM to 1 g wet biomass per 10 mL, and homogenised as above to yield a standardised inoculum.

### Time-lapse imaging of cuvette cultures

Macro cuvettes (Greiner Bio-One, Cat. no. 614101; 4 mL, 10 mm optical path; Greiner Bio-One, Kremsmünster, Austria) were each filled with 2 (species screen, and liquid level experiments) and 2.5 mL (membrane experiment) CYM and inoculated with 100 µL standardised inoculums. Cuvettes were held in a custom 3D-printed PLA rack positioned against the platen of an Epson Perfection V800 flatbed scanner (Seiko Epson Corporation, Suwa, Nagano, Japan) oriented vertically. Cultures were incubated at 27°C for the duration of each run. Scans were acquired hourly at 1,200 dpi (species screen and liquid level experiments) and 600 dpi (membrane experiment) in colour, scheduled with Cronicle, on a mini-PC running Manjaro Linux. Imaging was terminated after ∼ 10 days culturing. The five-species comparison used four replicate cuvettes per species (*n* = 4), and the cuvette liquid-level series used two to three replicates per level (*n* = 2–3).

### Image analysis

Images were processed in Python 3.13 with OpenCV 4.13. For each scan, a region of interest (ROI) was manually defined within the cuvette walls and slightly above the liquid-air interface to capture the pellicle formation. Mean grey intensity was computed per pixel row within the ROI to build spatiotemporal profiles, from which kymographs were assembled. Pixel-to-millimetre conversions were calculated by dividing the reference distance on the cuvette surface, measured with callipers (Mitutoyo Absolute AOS Digimatic digital), by its corresponding pixel length in the images.

Growth curves were obtained by averaging row-wise mean intensity across a cropped region around the liquid-air interface at each timepoint. A modified Gompertz function (18) was fitted to each curve by non-linear least-squares regression (SciPy curve_fit); lag-phase duration was taken directly as the fitted λ parameter, and time to stationary phase was defined as the time to reach 95% of the fitted intensity asymptote. Cultures of *Pleurotus ostreatus* never reached stationary phase during the experiment, and were thus excluded from this analysis.

### Pellicle harvesting and characterisation

Pellicles were lifted from cuvettes with a metal spatula. Adhering extracellular polymeric substance and submerged biomass were removed by gently scraping the underside under running tap water. Pellicles were laid on a silicone mat and air-dried under ambient conditions (25 °C, 30% relative humidity) for 3 d, then weighed on an analytical balance (Ohaus Adventurer, readability 0.1 mg). Thickness was measured with a digital thickness gauge (SHAHE, resolution 0.01 mm). Replicate numbers matched the corresponding culture experiments above. Pellicles from *P. ostreatus* were too fragile to handle and were thus excluded from biomass and thickness measurements.

### Membrane-separated culturing

Cultures were prepared and imaged as for the cuvette experiments above, except that the working volume was increased to 2.5 mL while maintaining a 100 µL inoculum. A custom 3D-printed lid bearing a 0.22 µm cellulose acetate membrane (Ji’an City Qing Feng Filter Equipment Material Co., Ltd, Jiangxi, China) was fitted to each cuvette, with the membrane bonded to the lid using cyanoacrylate adhesive. Assemblies were sterilised by immersion in 70% ethanol for 1 h followed by UV irradiation for 30 min before insertion into cuvettes. The membrane retained inoculum at the liquid–air interface while permitting nutrient transport. The experiment compared membrane and no-membrane conditions with ten replicates each (n = 10).

### Statistics and reproducibility

All statistical analyses were performed in Python 3.13.2 using Pingouin 0.6.0, SciPy 1.17.1, scikit-learn 1.8.0, statsmodels 0.14.6, PyMC 5.28.1 and ArviZ 0.23.4, with NumPy 2.4.3 and pandas 3.0.1. Replicate numbers are biologically independent cultures and are stated for each experiment in the relevant sections. All frequentist tests were two-sided with significance assessed at α = 0.05; figure error bars represent standard deviations.

Growth parameters were obtained by fitting a Gompertz–Zwietering model to each ROI-averaged growth curve by non-linear least-squares regression (scipy.optimize.curve_fit). Lag-phase duration was taken directly as the fitted λ parameter, and time to stationary phase was defined as the time at which the fitted curve first reached 95% of its amplitude (by linear interpolation). Uncertainty in the fitted parameters and in the derived stationary time was estimated by residual bootstrap (B = 300 resamples; percentile 95% confidence intervals), using a seeded random generator so that the reported intervals are reproducible.

For between-group comparisons, homogeneity of variance was first tested by the Brown– Forsythe test (median-centred Levene). When variances were homogeneous, groups were compared by one-way ANOVA with Tukey HSD post hoc tests; when variances were heterogeneous, Welch’s ANOVA with Games–Howell post hoc tests were used instead. On this basis, lag phase, biomass yield and pellicle thickness were analysed by one-way ANOVA with Tukey HSD, whereas time to stationary phase was analysed by Welch’s ANOVA with Games–Howell. Omnibus effect sizes are reported as partial η², and pairwise effect sizes as small-sample-corrected Hedges’ g. For the two-condition membrane experiment, lag phase, time to stationary phase and biomass were compared by Welch’s ANOVA (equivalent to a Welch’s t-test for two groups) with Games–Howell tests and Hedges’ g, and equality of variances between conditions was assessed using both the Brown–Forsythe and Fligner–Killeen tests.

The dependence of growth parameters on liquid depth was characterised by fitting candidate dose–response models (linear, logarithmic, and a saturating Michaelis– Menten-type function) by non-linear least-squares regression, with competing models compared by the difference in Akaike information criterion (ΔAIC) and the strength of monotonic association quantified by Spearman’s rank correlation (ρ). Heteroscedasticity of responses with respect to liquid depth were tested by ordinary least-squares regression followed by the Breusch–Pagan and Goldfeld–Quandt tests.

To separate biological variability from measurement uncertainty in lag time, a Bayesian heteroscedastic regression was fitted in PyMC. Replicate lag times were modelled as y ∼ Normal(α + βz, σ_tot_), where z is the standardised liquid depth (log-transformed for time to stationary phase), the biological standard deviation σ_bio_ = exp(γ₀ + γ₁z) varies with depth, and σ_tot_ = 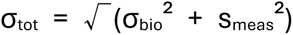 incorporates the per-replicate measurement uncertainty s_meas (the bootstrap lag-time SD). Priors were α ∼ Normal(ȳ, 10), β ∼ Normal(0, 10), γ₀ ∼ Normal(ln SD(y), 2) and γ₁ ∼ Normal(0, 1). Posteriors were sampled using the No-U-Turn sampler (4 chains; 2,000 tuning and 2,000 post-tuning draws per chain; target acceptance 0.9); convergence was confirmed with all R̂ ≈ 1.00 and adequate effective sample sizes. Posterior estimates are summarised as the posterior mean and 94% highest-density interval (HDI), and the hypothesis that biological variability increases with liquid depth was quantified as the posterior probability that γ₁ > 0.

## Data availability

All supporting data has been provided in the Supplementary Materials. Raw data, and analysis code has been provided as an open-source repository. See DOI: (10.5281/zenodo.22656014).

## CRediT authorship contribution statement

Piet Dimitriadis: Conceptualization, Data curation, Formal Analysis, Investigation, Methodology, Project administration, Software, Visualization, Writing – original draft, Writing – review & editing; Anouk Verstuyft: Visualization, Writing – review & editing; Jara Saluena Martin: Conceptualization, Writing – review & editing; Rafael Philippini: Conceptualization, Writing – review & editing; Annah-Ololade Sangosanya: Writing – review & editing; Simon Vandelook: Conceptualization, Writing – review & editing; Wim de Malsche: Conceptualization, Supervision, Writing – review & editing; Eveline Peeters: Conceptualization, Project administration, Resources, Supervision, Writing – review & editing

