## Supplementary Materials for "Inoculum-to-interface distance tunes productivity in static liquid cultures of wood-decay fungi"

Dimitriadis *et al.*

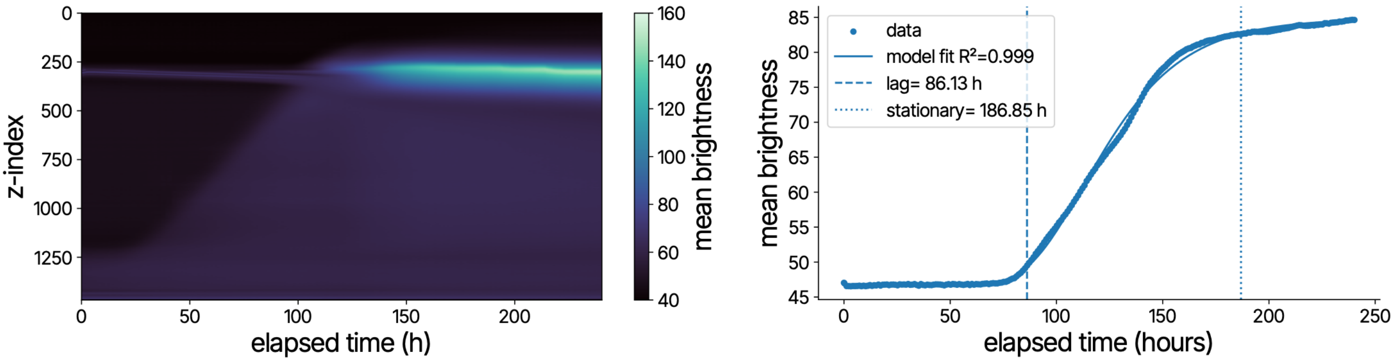


***Fig S1. Kymograph and growth curve of ROI 0 containing T. versicolor***

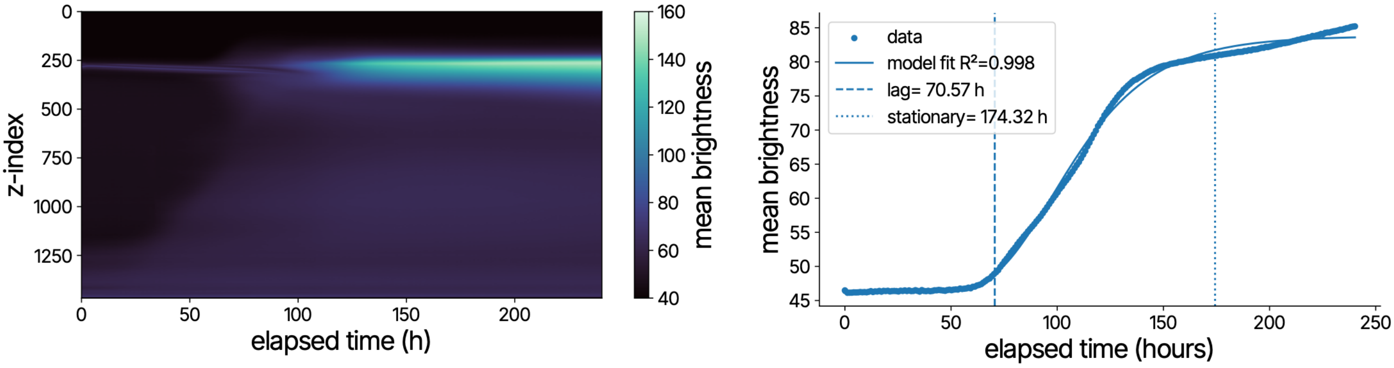


***Fig S2. Kymograph and growth curve of ROI 1 containing T. versicolor***

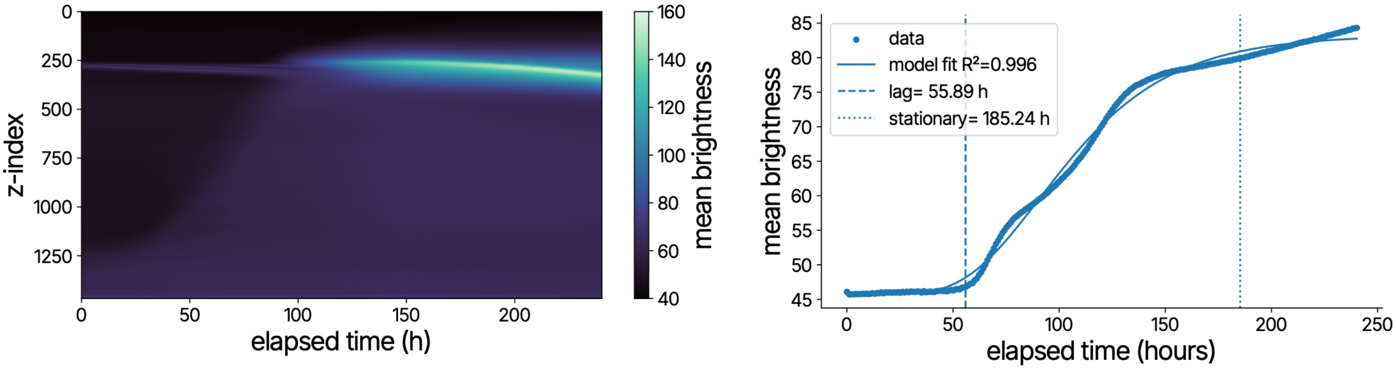


***Fig S3. Kymograph and growth curve of ROI 2 containing T. versicolor***

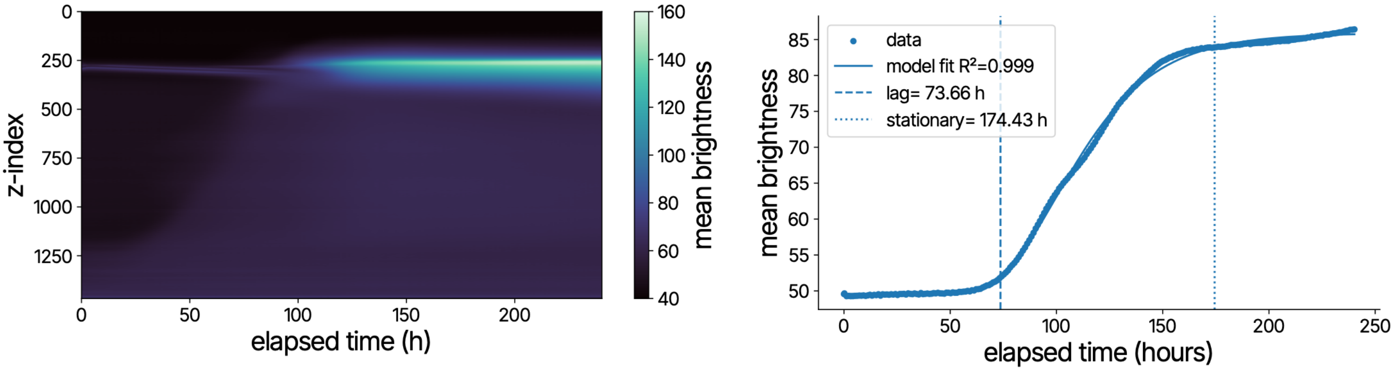


*Fig S4. Kymograph and growth curve of ROI 3 containing T. versicolor*

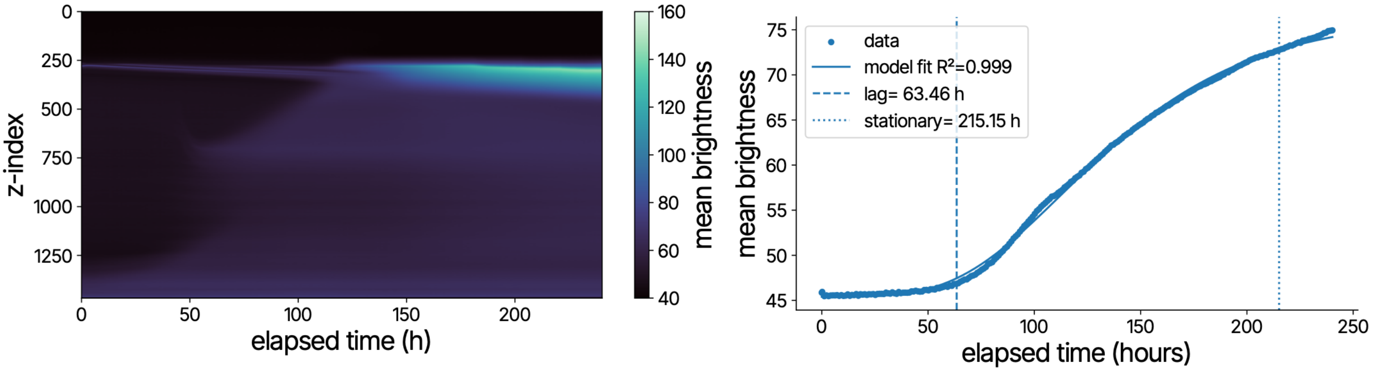


*Fig S5. Kymograph and growth curve of ROI 4 containing G. sessile*

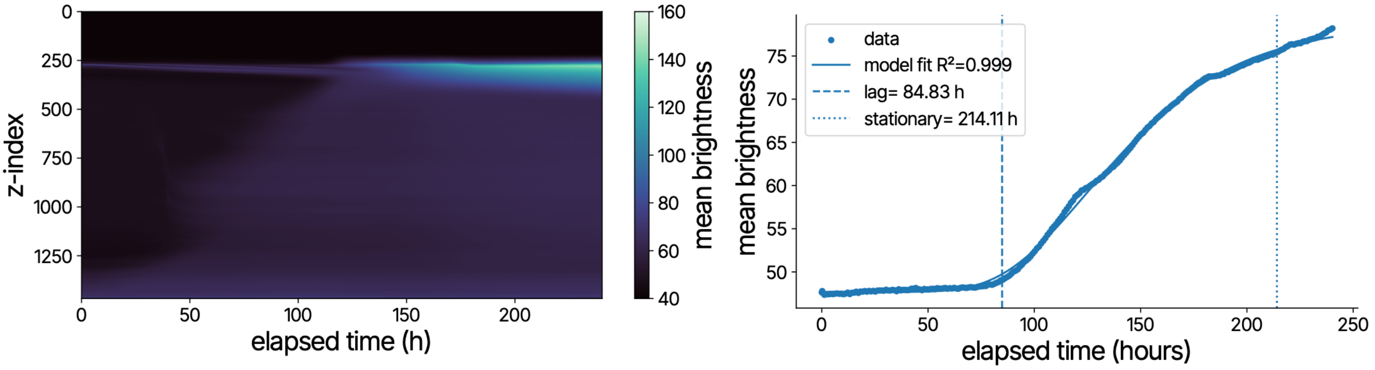


*Fig S6. Kymograph and growth curve of ROI 5 containing G. sessile*

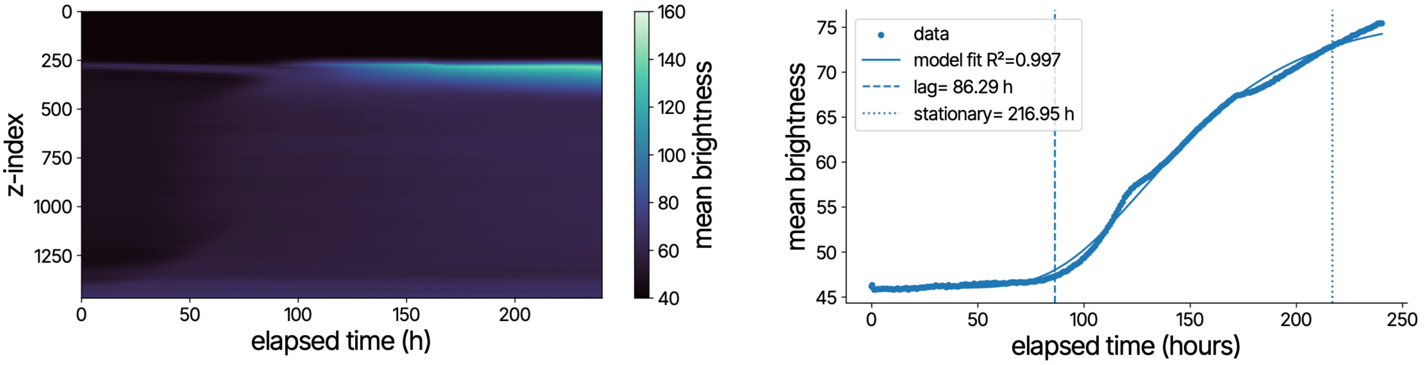


*Fig S7. Kymograph and growth curve of ROI 6 containing G. sessile*

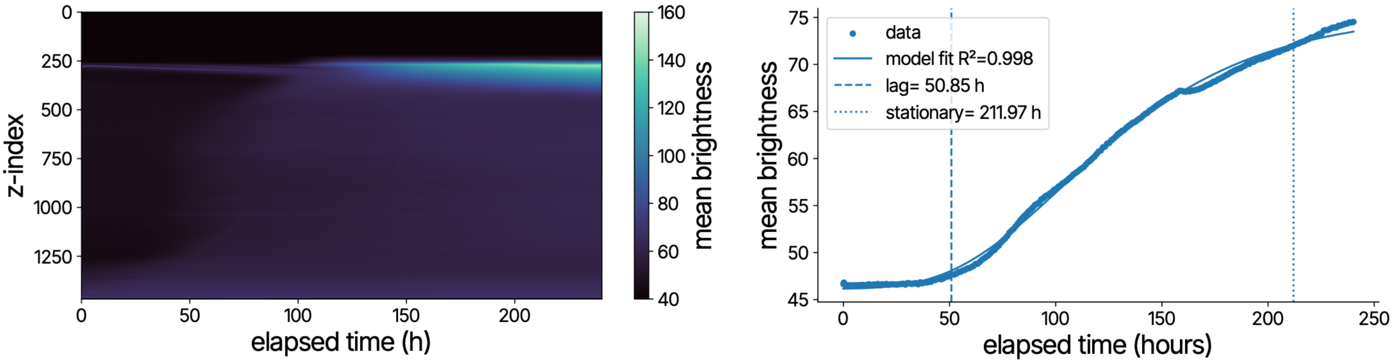


*Fig S8. Kymograph and growth curve of ROI 7 containing G. sessile*

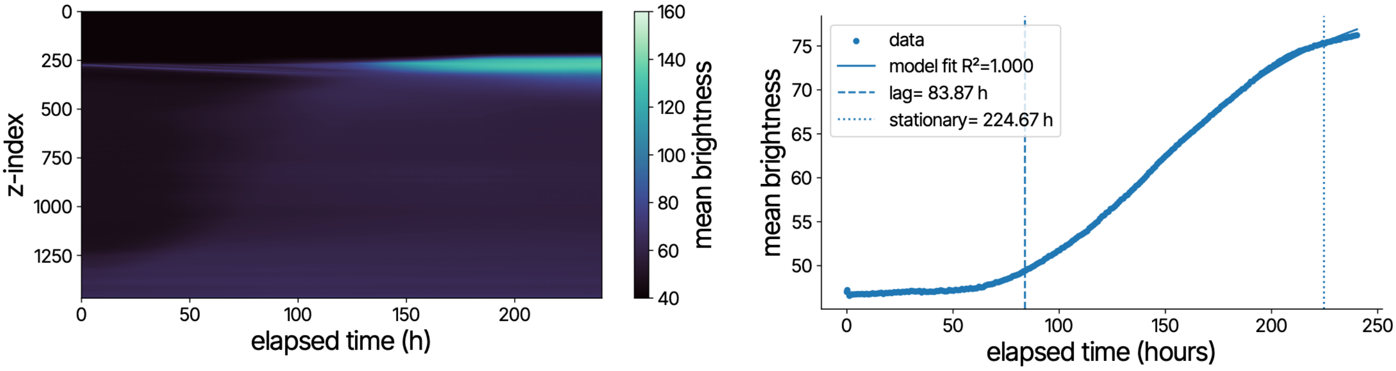


*Fig S9. Kymograph and growth curve of ROI 8 containing F. fomentarius*

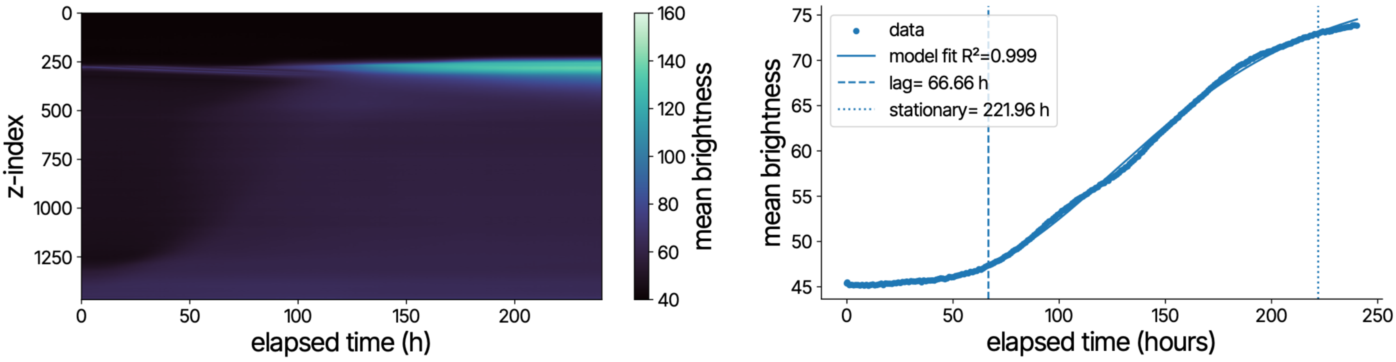


*Fig S10. Kymograph and growth curve of ROI 9 containing F. fomentarius*

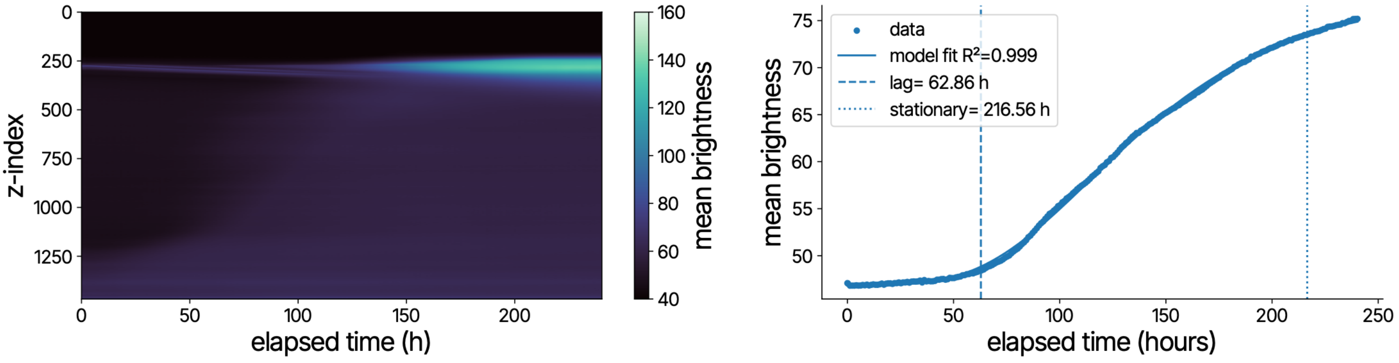


*Fig S11. Kymograph and growth curve of ROI 10 containing F. fomentarius*

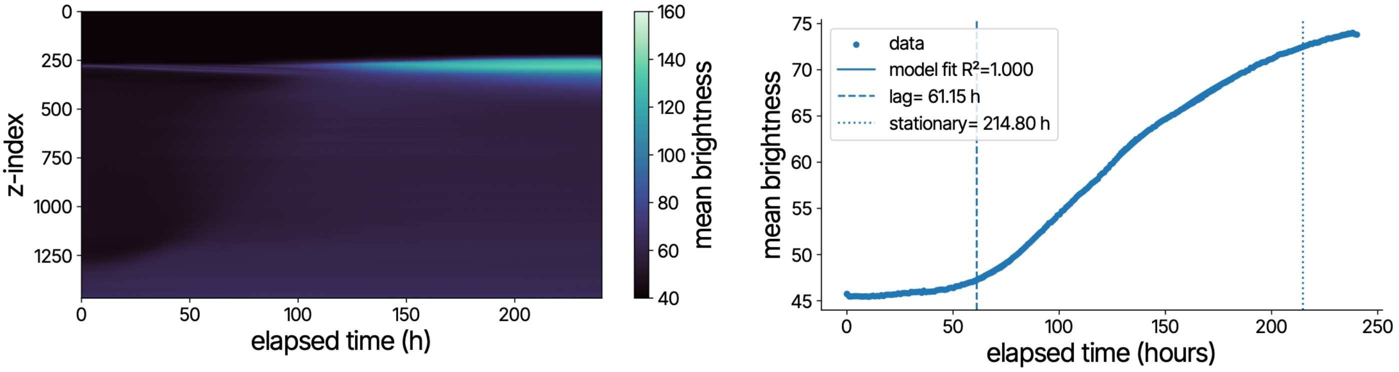


*Fig S12. Kymograph and growth curve of ROI 11 containing F. fomentarius*

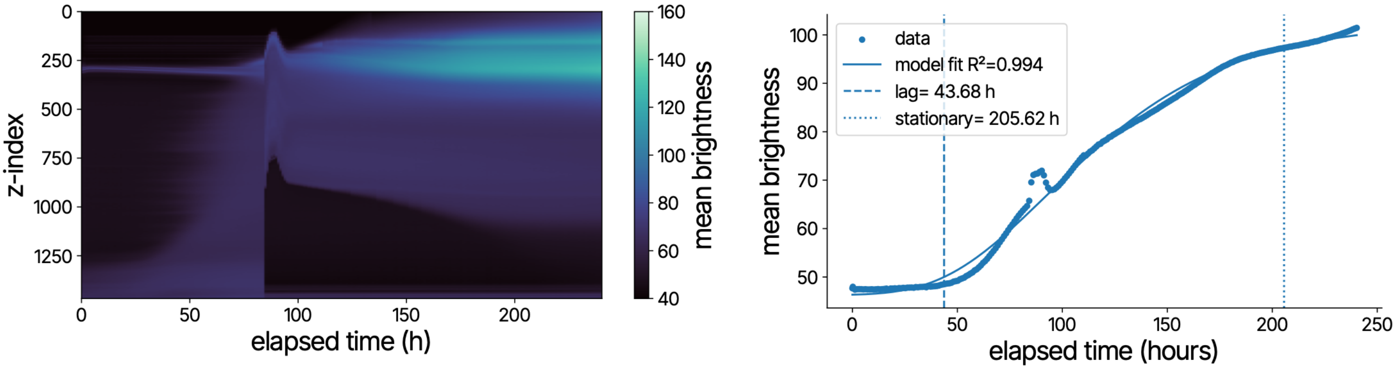


*Fig S13. Kymograph and growth curve of ROI 12 containing S. commune*

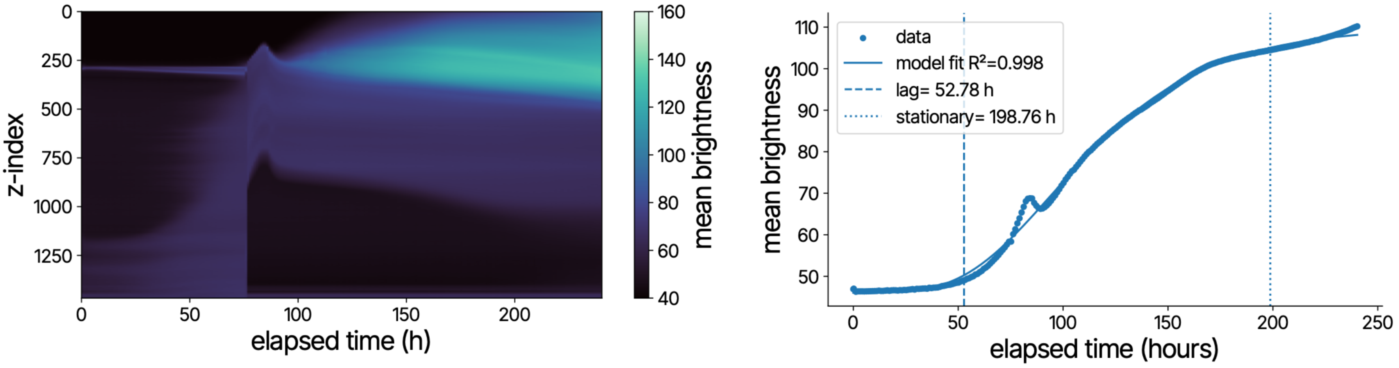


*Fig S14. Kymograph and growth curve of ROI 13 containing S. commune*

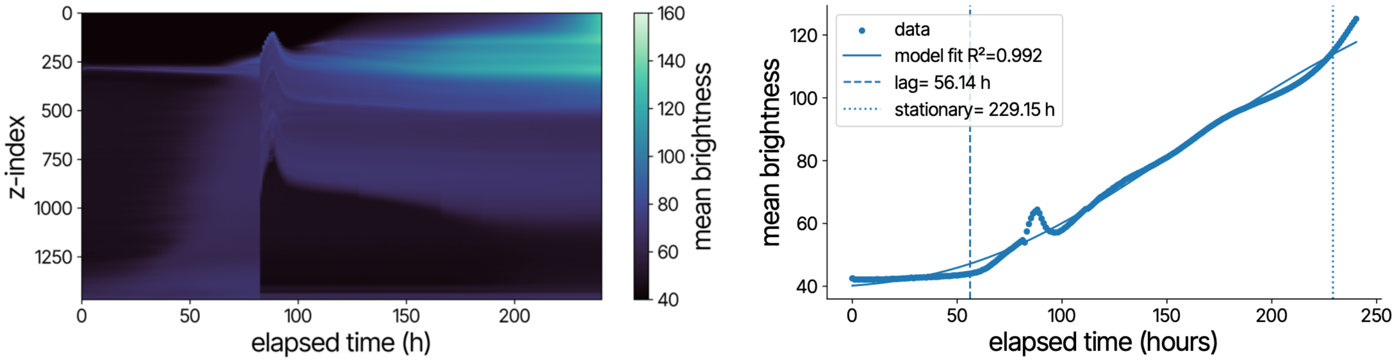


*Fig S15. Kymograph and growth curve of ROI 14 containing S. commune*

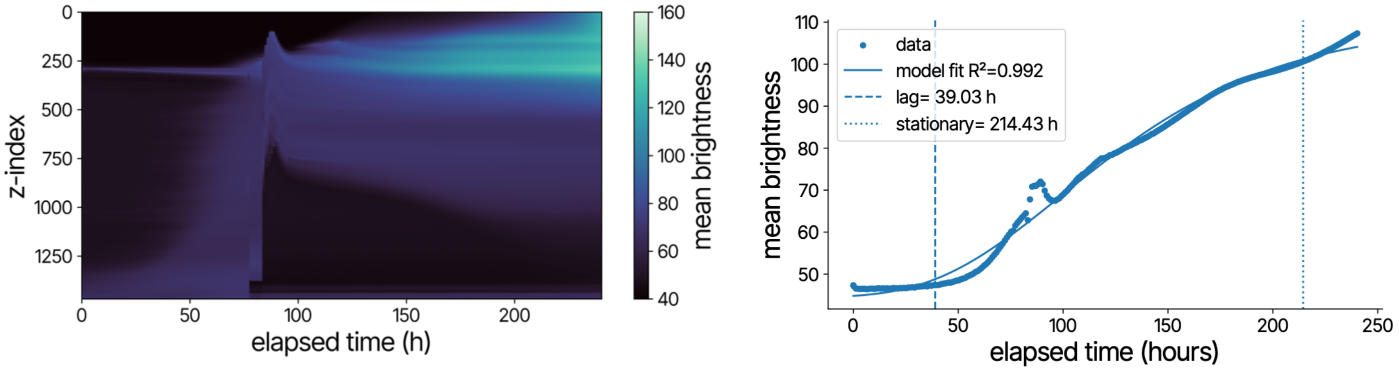


*Fig S16. Kymograph and growth curve of ROI 15 containing S. commune*

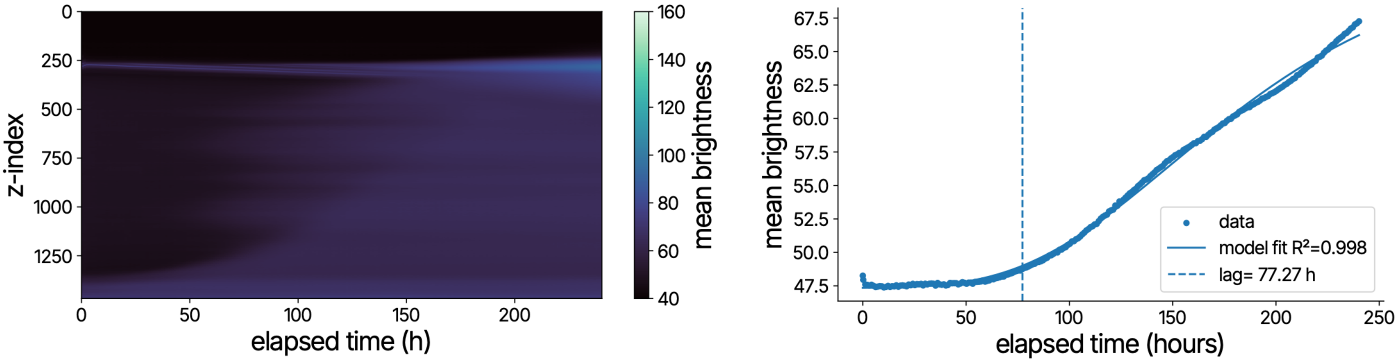


*Fig S17. Kymograph and growth curve of ROI 16 containing P. ostreatus*

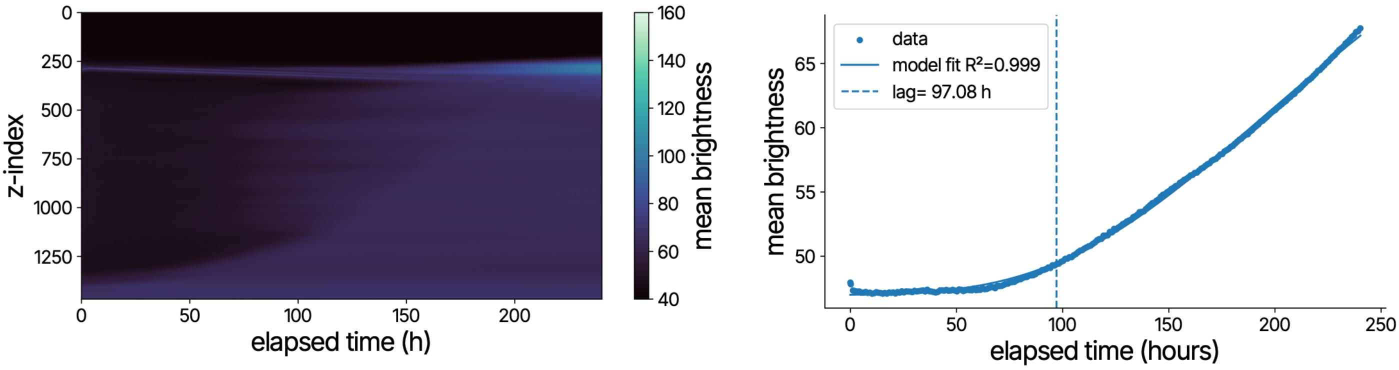


*Fig S18. Kymograph and growth curve of ROI 17 containing P. ostreatus*

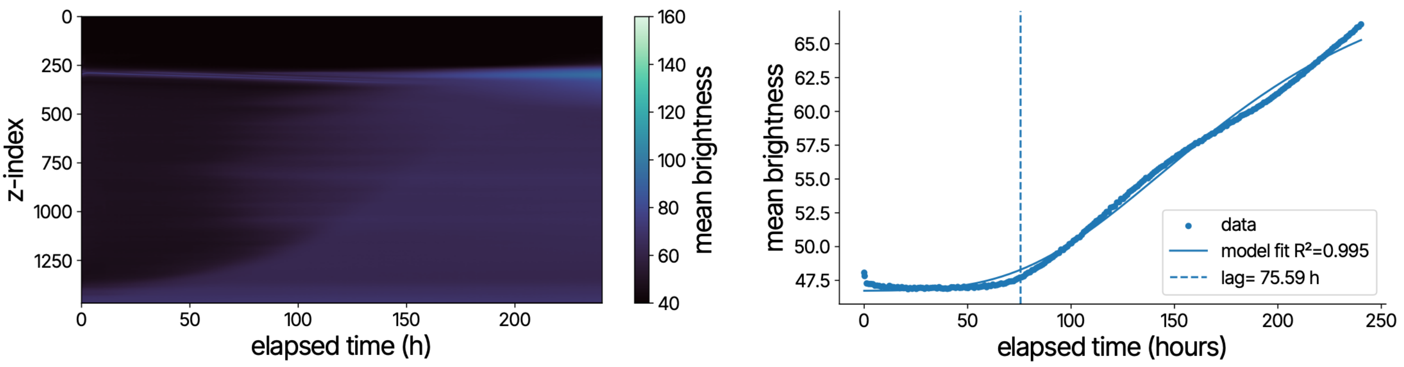


***Fig S19. Kymograph and growth curve of ROI 18 containing P. ostreatus***

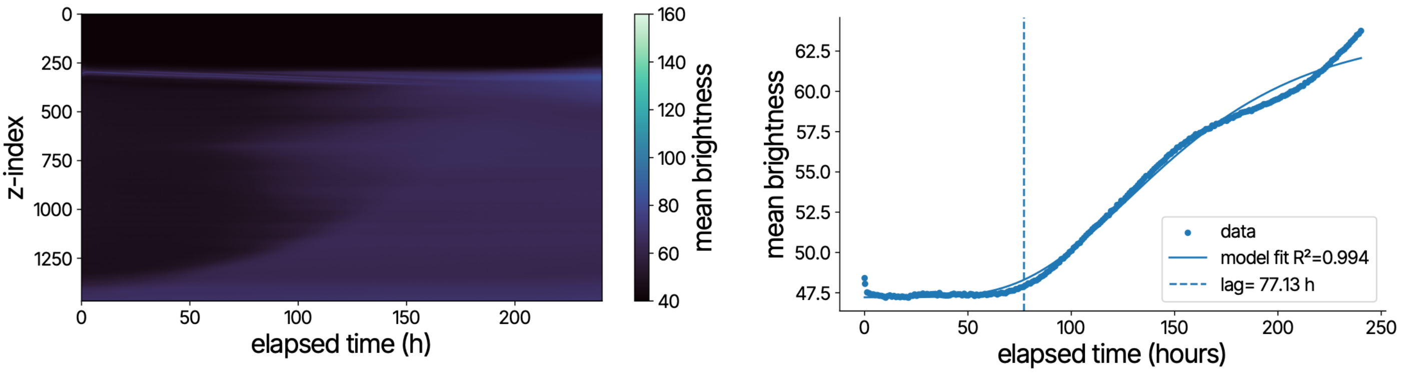


***Fig S20. Kymograph and growth curve of ROI 19 containing P. ostreatus***

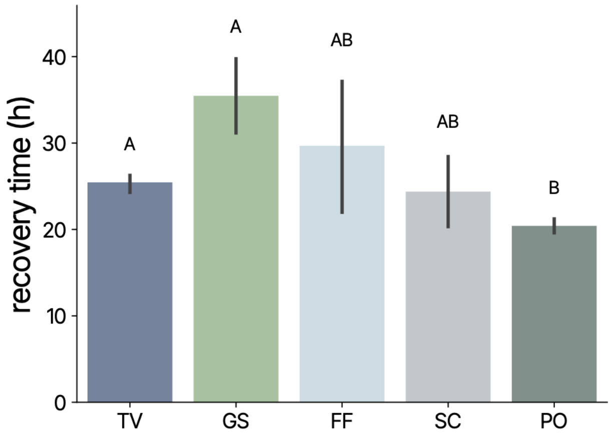


***Fig S21 Species selection influences recovery time.*** *Groups sharing a letter in the compact-letter display are not distinguishable at α = 0.05 (Welch's ANOVA).*

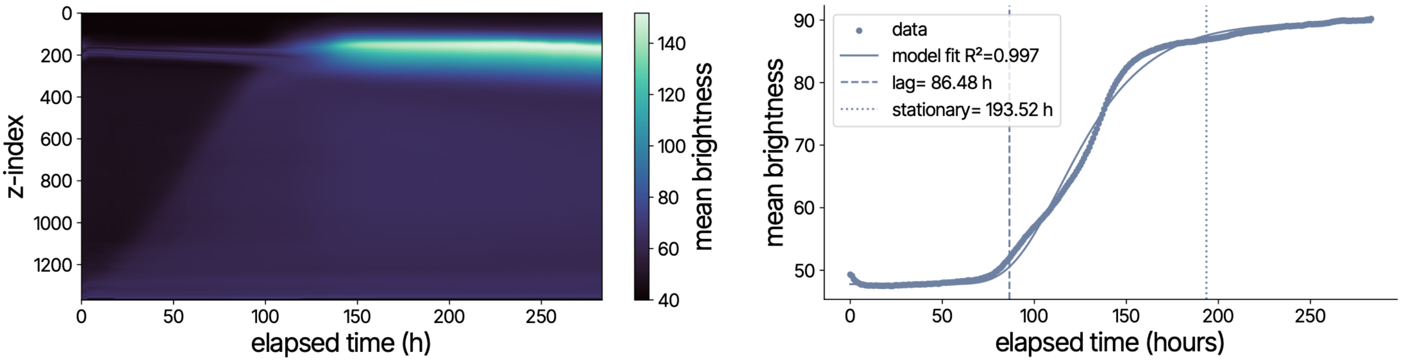


*Fig S22. Kymograph and growth curve of ROI 0 at Δz = 20.7 mm*

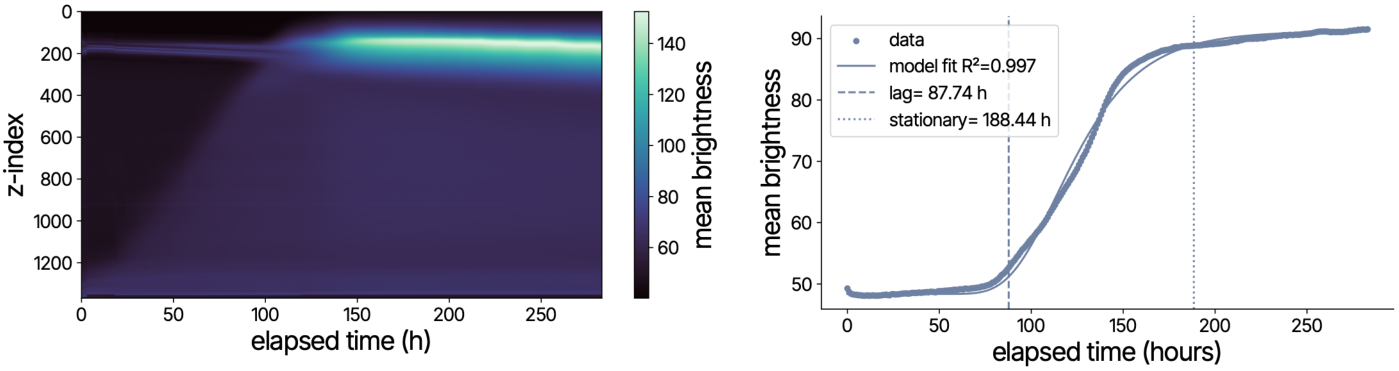


*Fig S23. Kymograph and growth curve of ROI 1 at Δz = 20.7 mm*

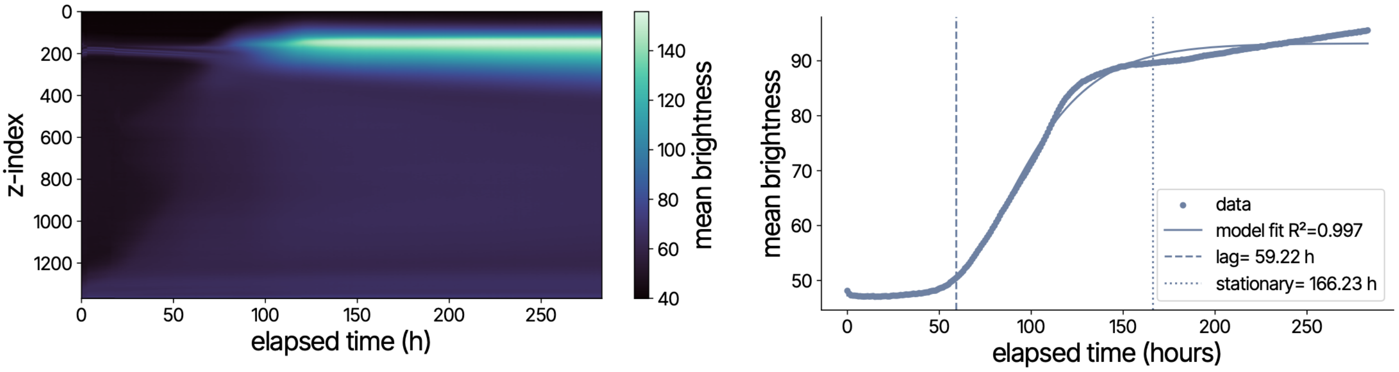


*Fig S24. Kymograph and growth curve of ROI 2 at Δz = 20.7 mm*

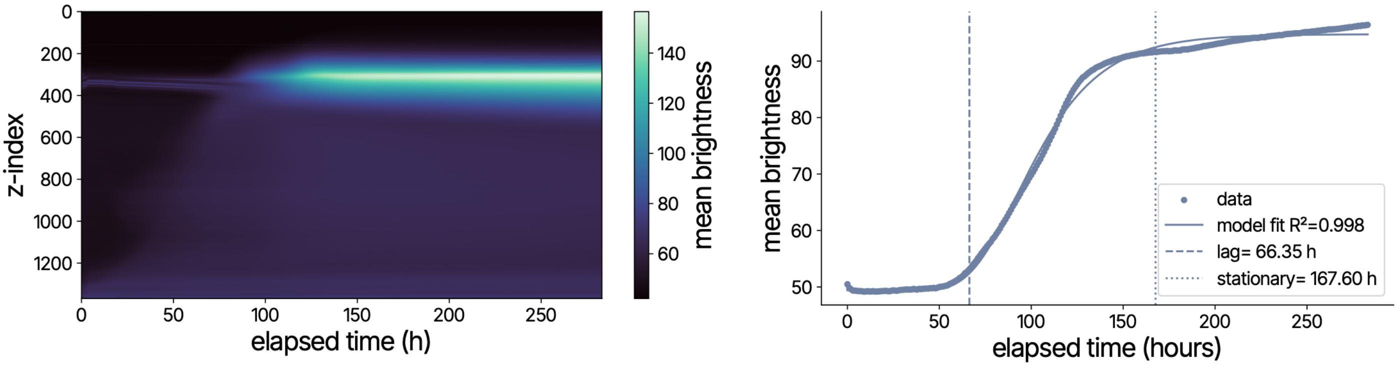


*Fig S25. Kymograph and growth curve of ROI 3 at Δz = 17.5 mm*

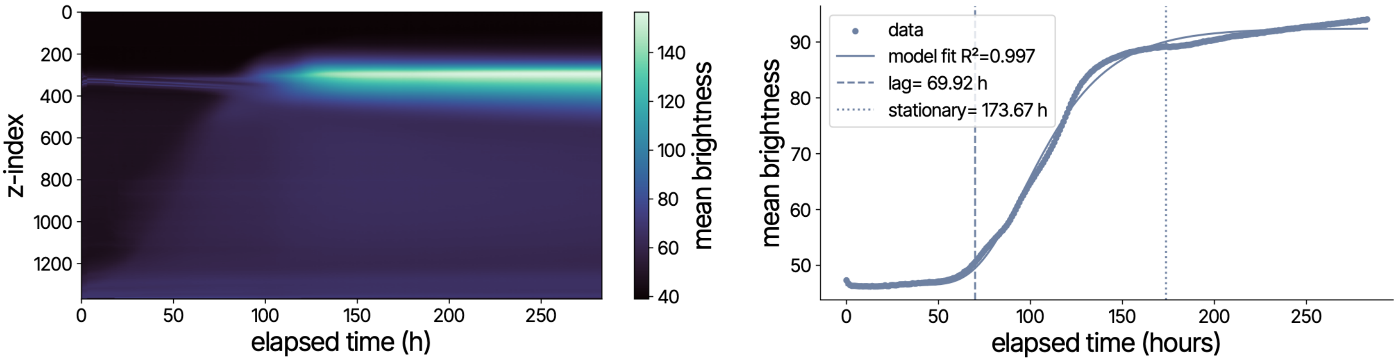


*Fig S26. Kymograph and growth curve of ROI 4 at Δz = 17.5 mm*

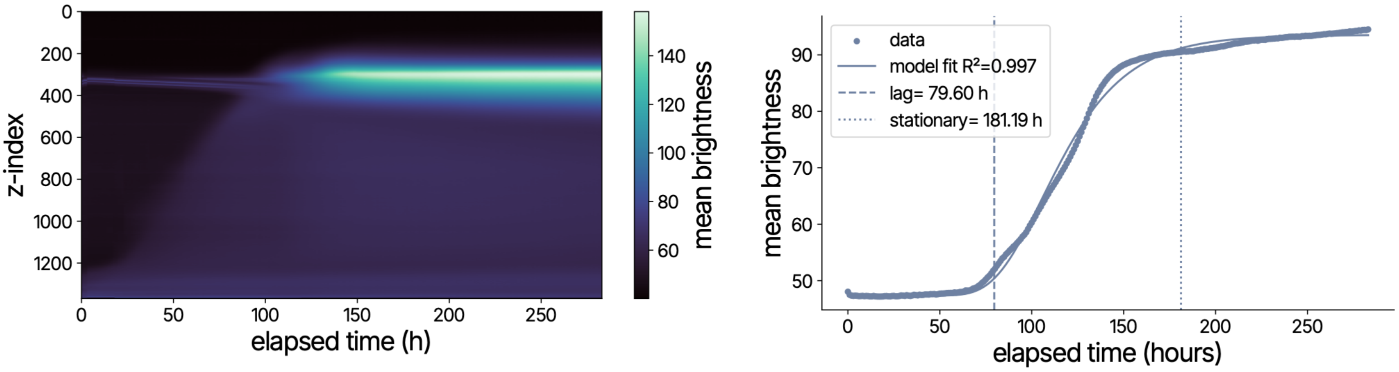


*Fig S27. Kymograph and growth curve of ROI 5 at Δz = 17.5 mm*

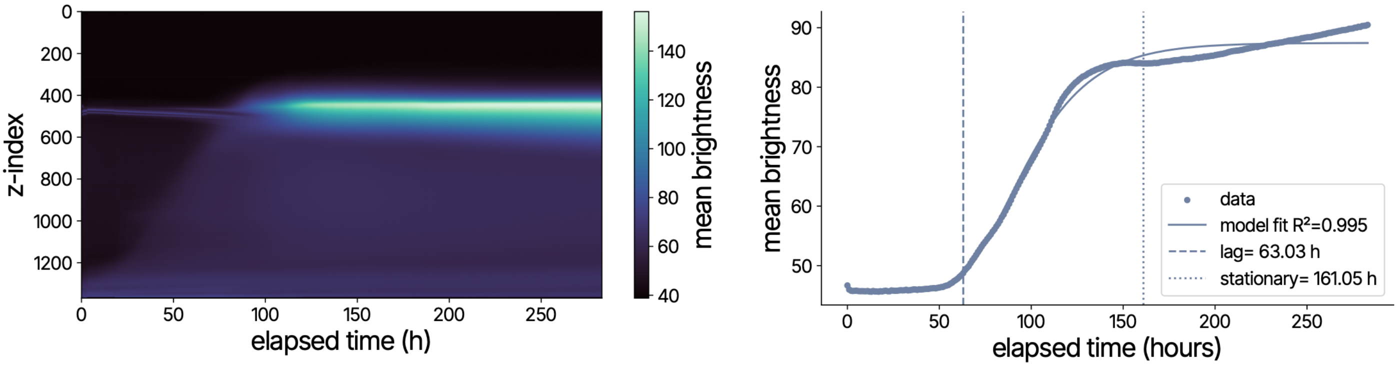


*Fig S28. Kymograph and growth curve of ROI 6 at Δz = 14.0 mm*

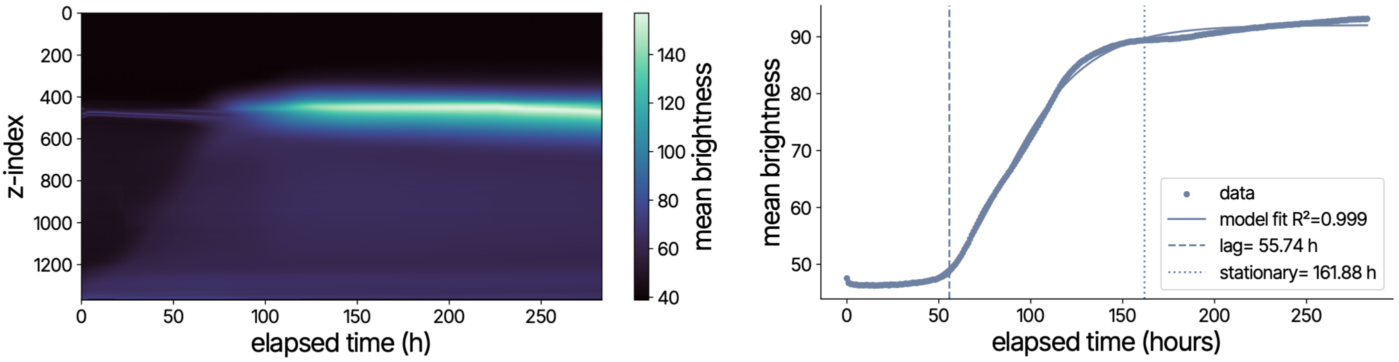


*Fig S29. Kymograph and growth curve of ROI 7 at Δz = 14.0 mm*

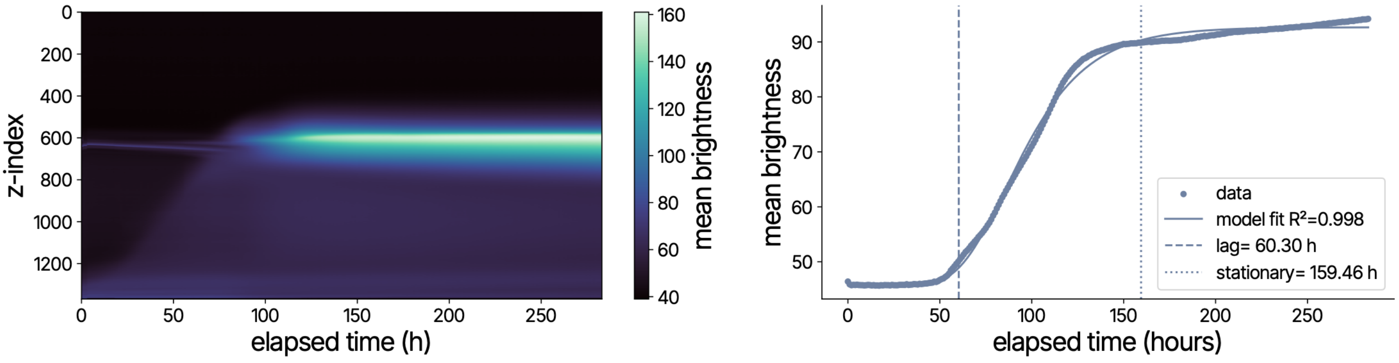


*Fig S30. Kymograph and growth curve of ROI 8 at Δz = 10.68 mm*

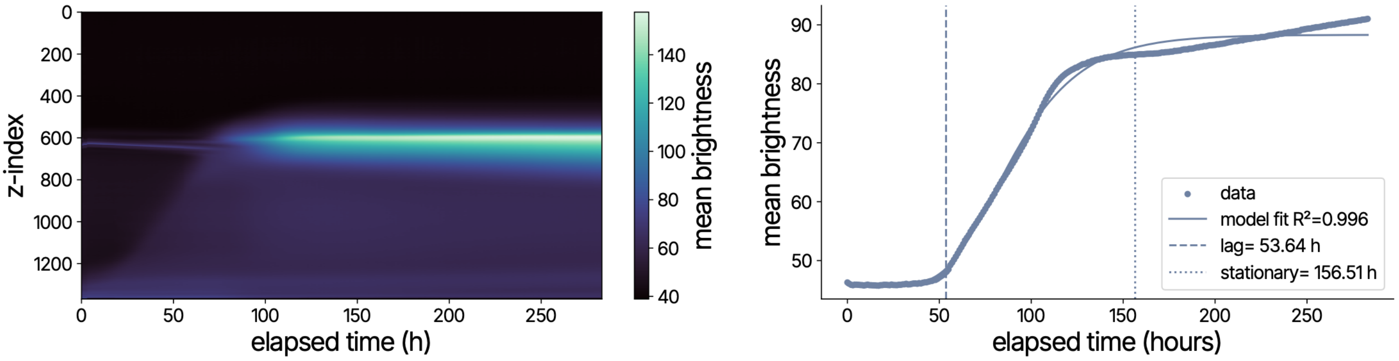


*Fig S31. Kymograph and growth curve of ROI 9 at Δz = 10.68 mm*

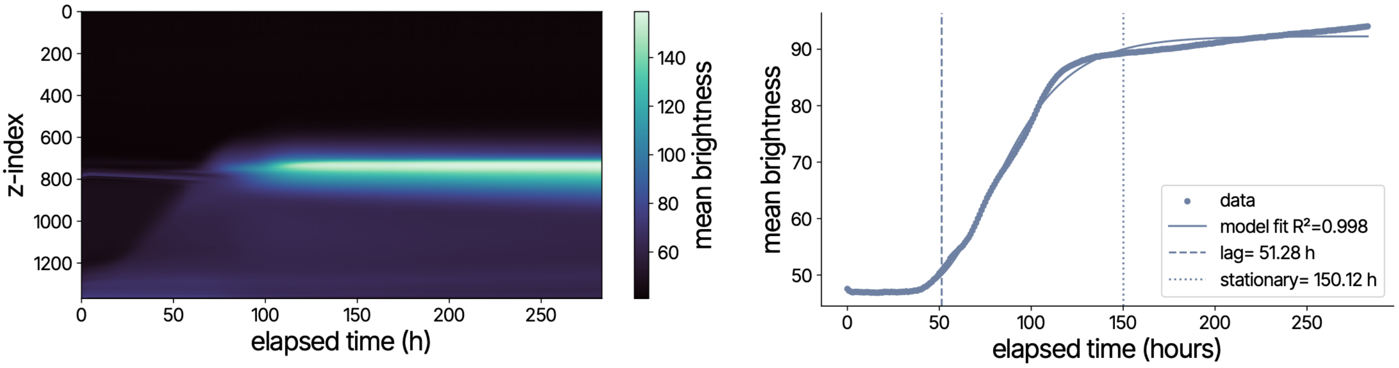


*Fig S32. Kymograph and growth curve of ROI 10 at Δz = 7.65 mm*

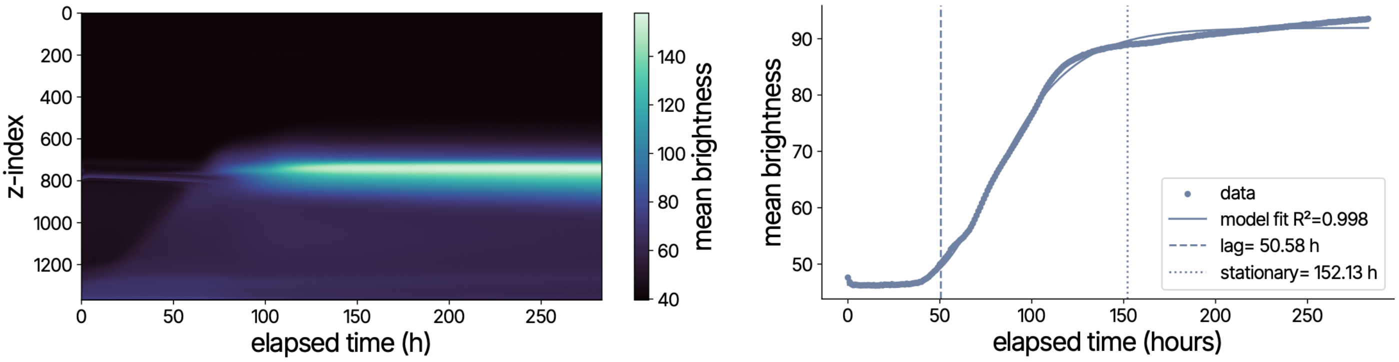


*Fig S33. Kymograph and growth curve of ROI 11 at Δz = 7.65 mm*

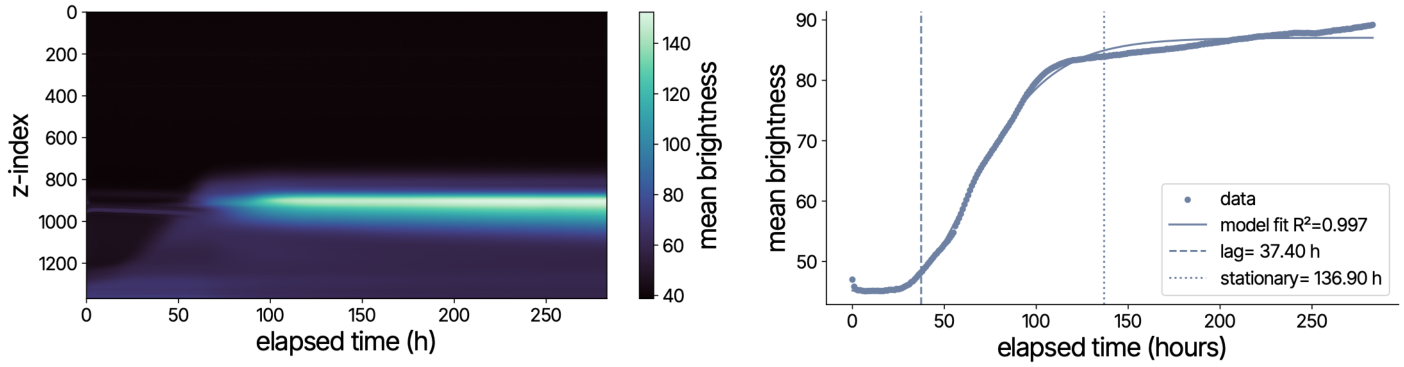


*Fig S34. Kymograph and growth curve of ROI 12 at Δz = 4.92 mm*

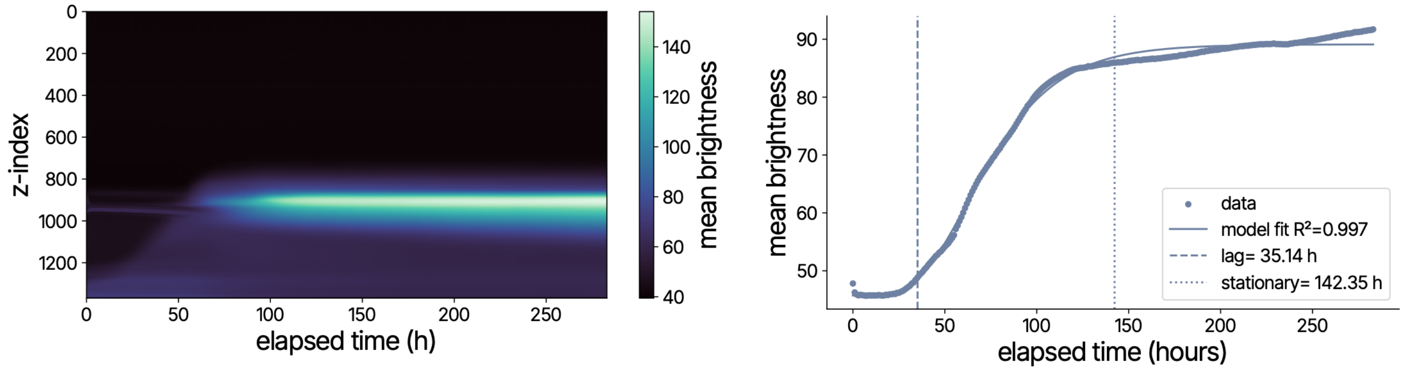


*Fig S35. Kymograph and growth curve of ROI 13 at Δz = 4.92 mm*

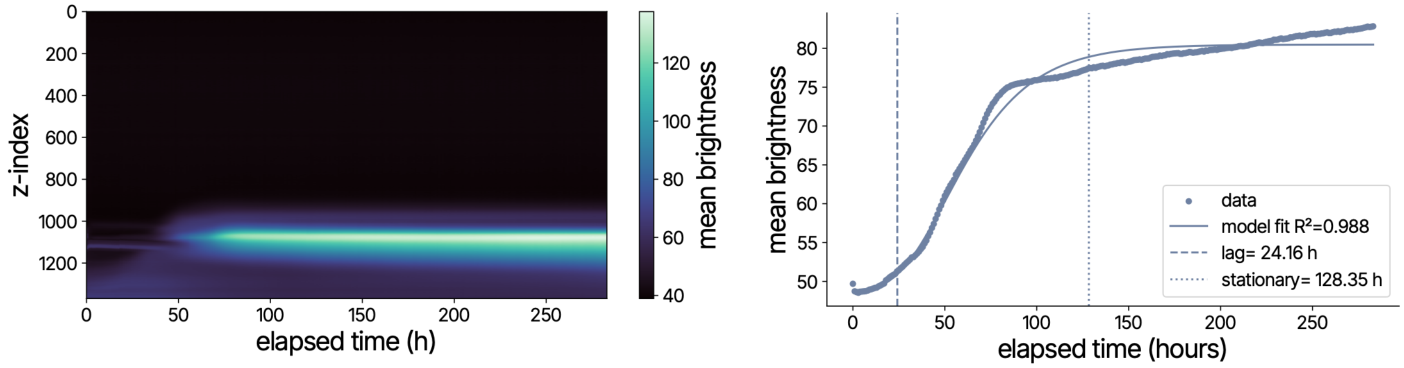


*Fig S36. Kymograph and growth curve of ROI 14 at Δz = 1.33 mm*

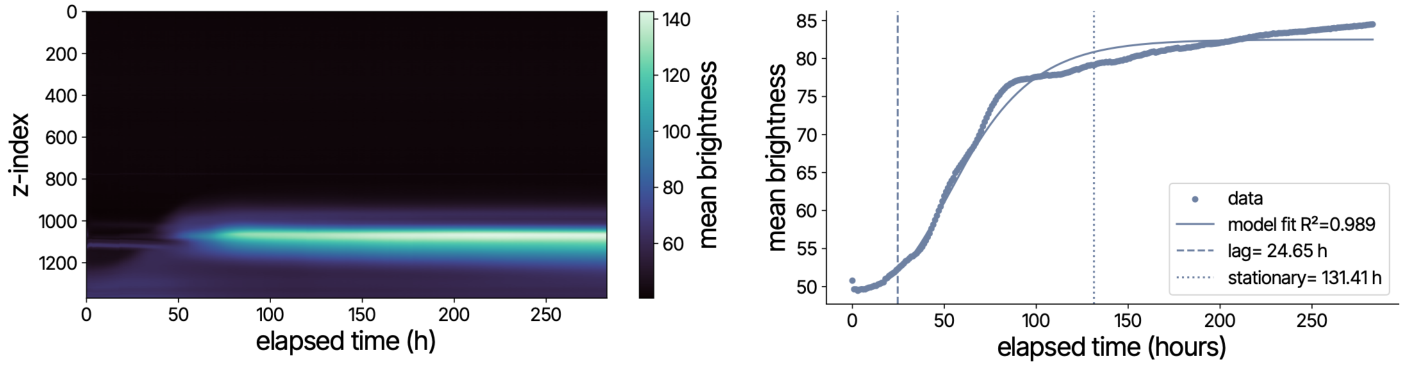


*Fig S37. Kymograph and growth curve of ROI 15 at Δz = 1.33 mm*

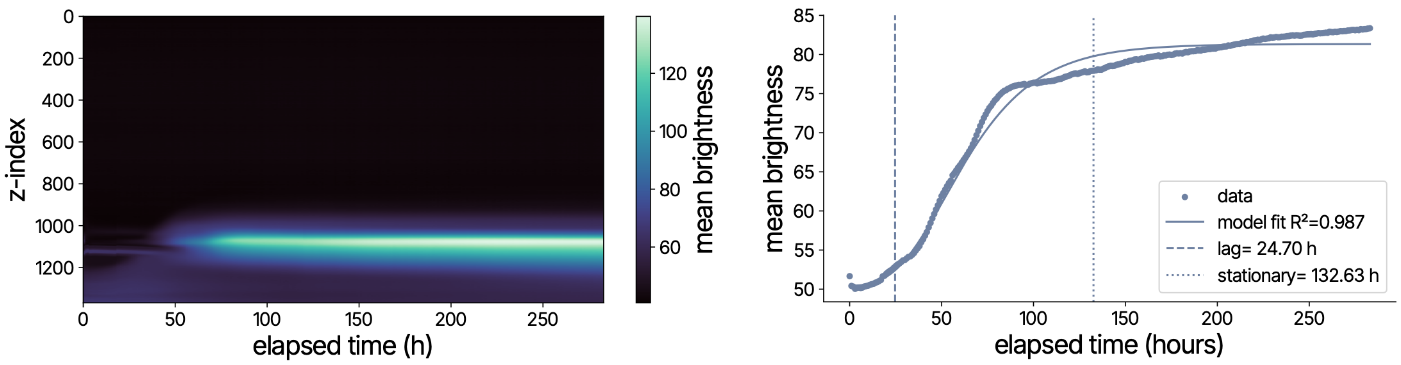


*Fig S38. Kymograph and growth curve of ROI 16 at Δz = 1.33 mm*

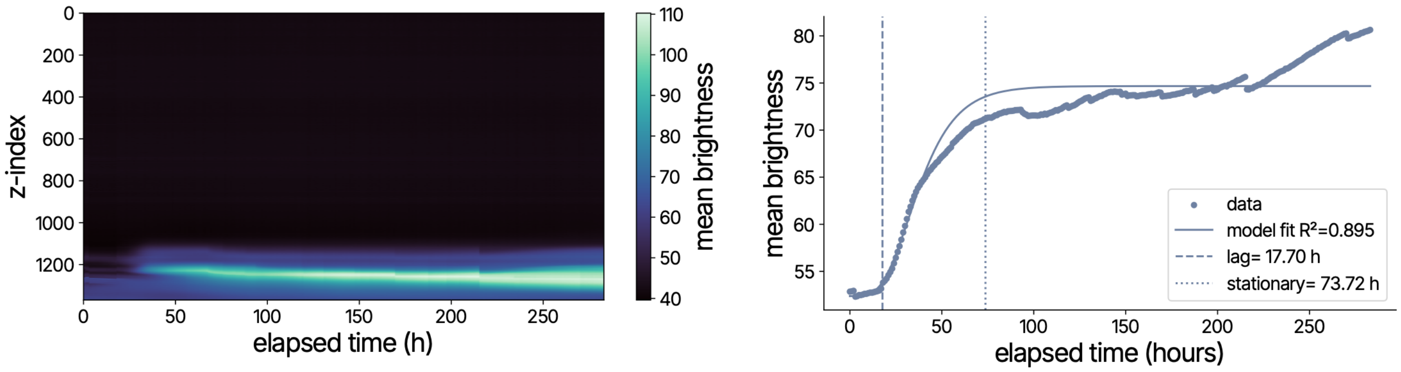


*Fig S39. Kymograph and growth curve of ROI 17 at Δz = 0.45 mm*

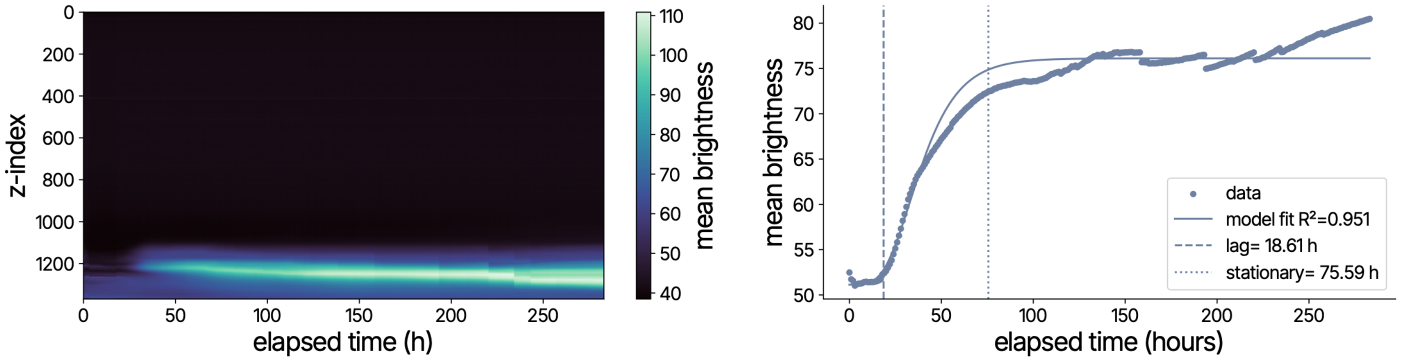


*Fig S40. Kymograph and growth curve of ROI 18 at Δz = 0.45 mm*

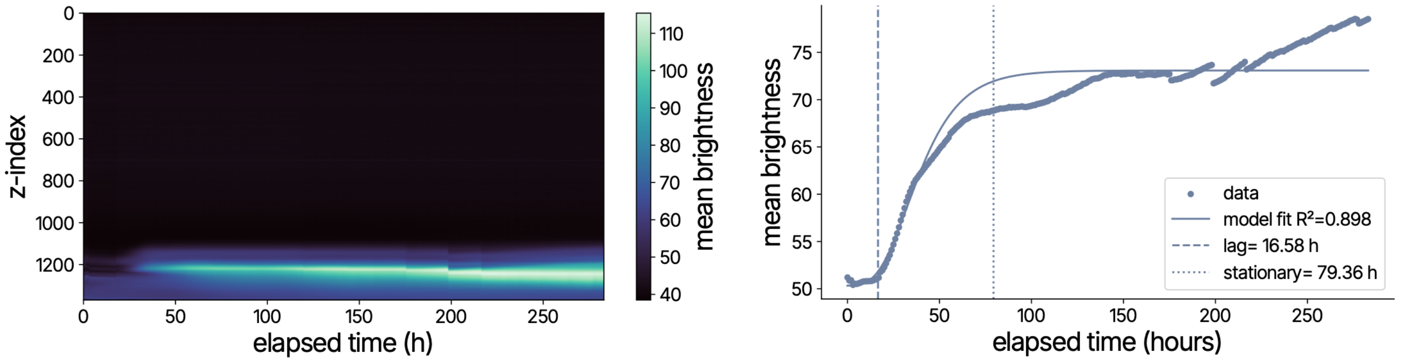


*Fig S41. Kymograph and growth curve of ROI 19 at Δz = 0.45 mm*

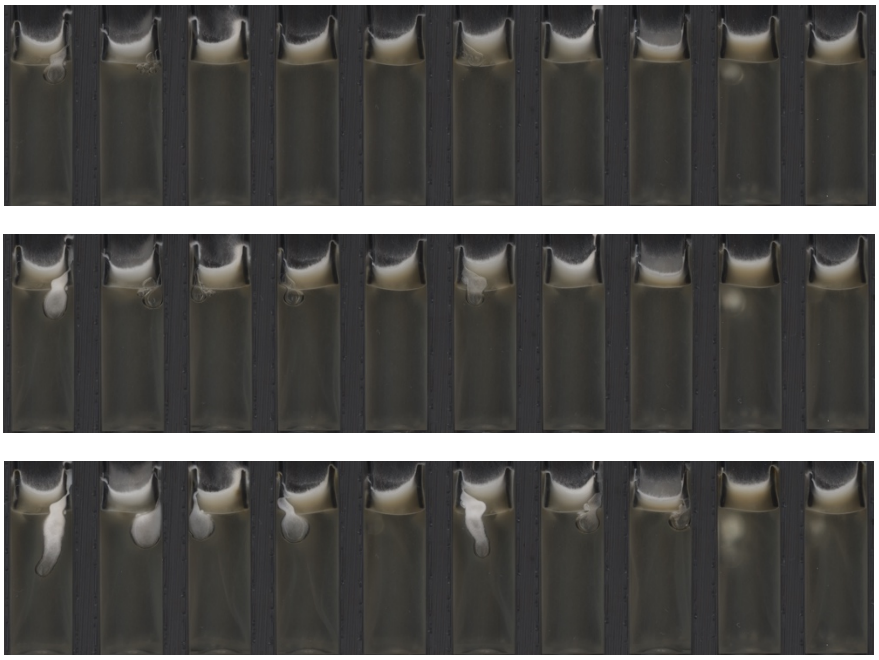

*Fig S42. Hyphae colonise gas pockets trapped below the membrane, increasing pellicle biomass*

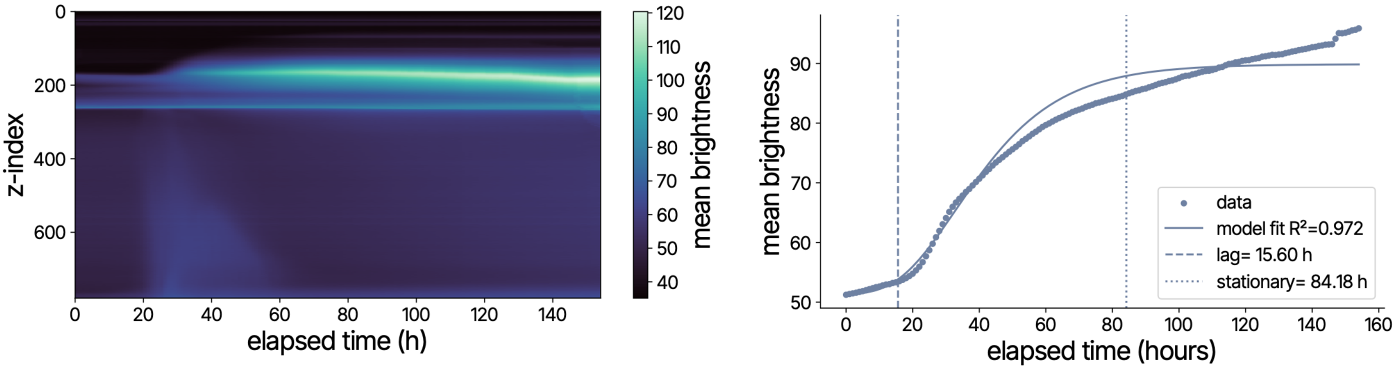


*Fig S43. Kymograph and growth curve of ROI 0 with membrane*

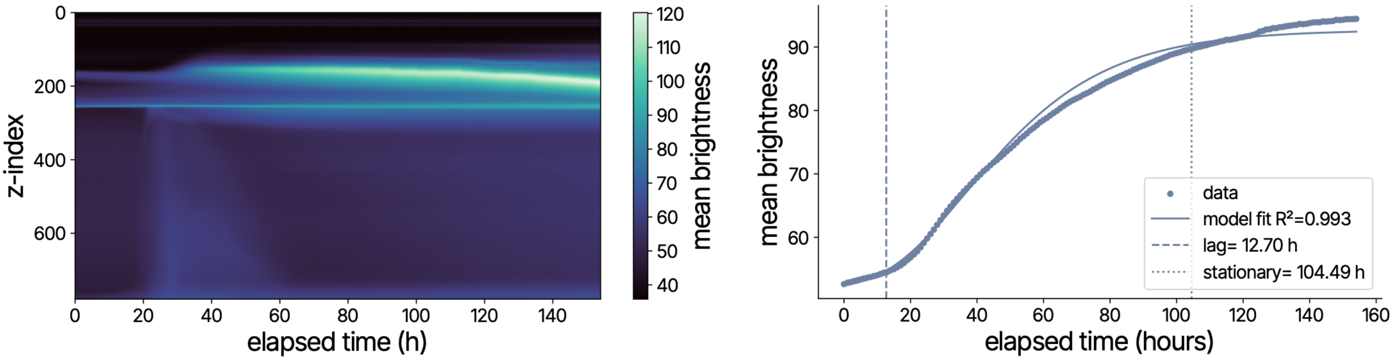


*Fig S44. Kymograph and growth curve of ROI 1 with membrane*

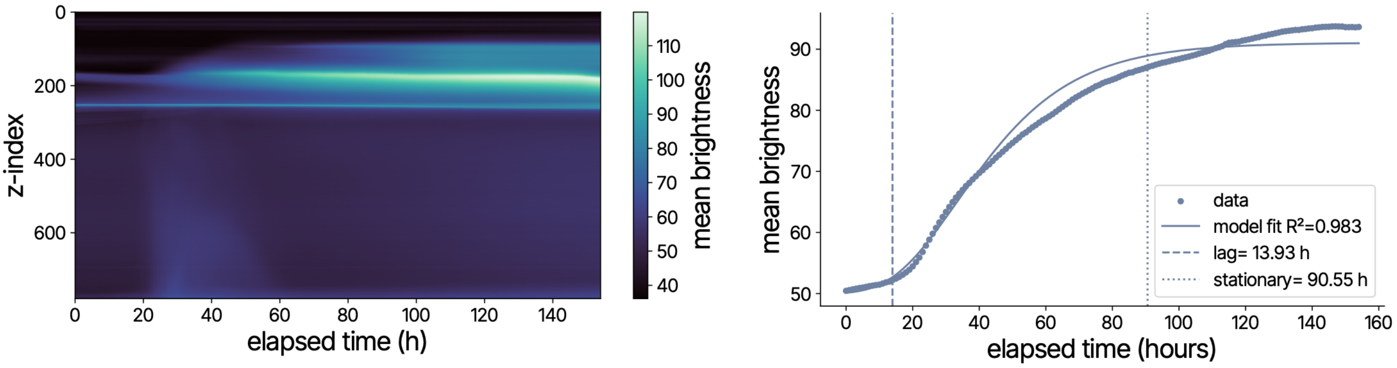


*Fig S45. Kymograph and growth curve of ROI 2 with membrane*

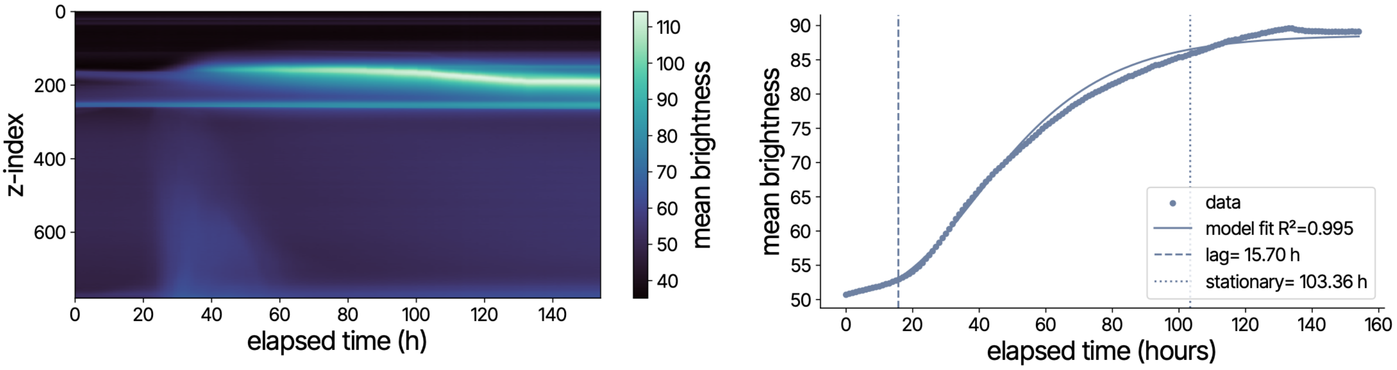


*Fig S46. Kymograph and growth curve of ROI 3 with membrane*

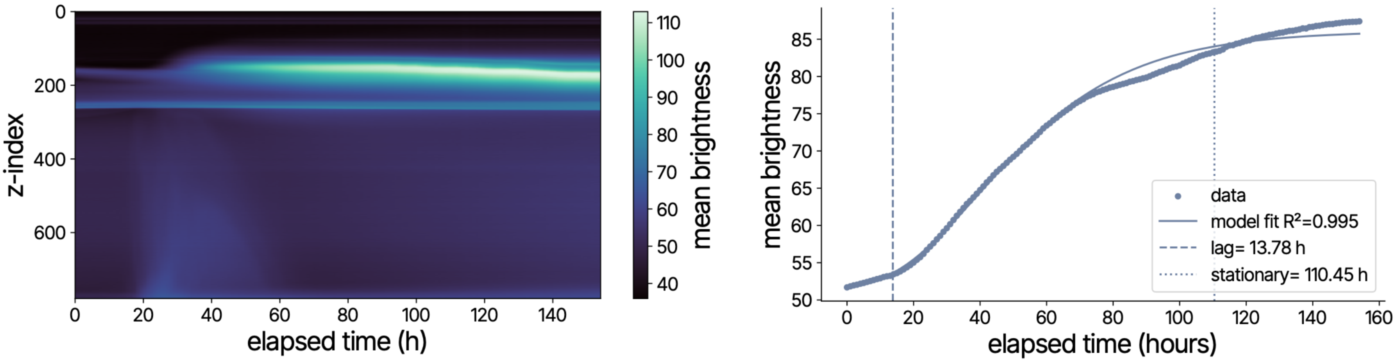


*Fig S47. Kymograph and growth curve of ROI 4 with membrane*

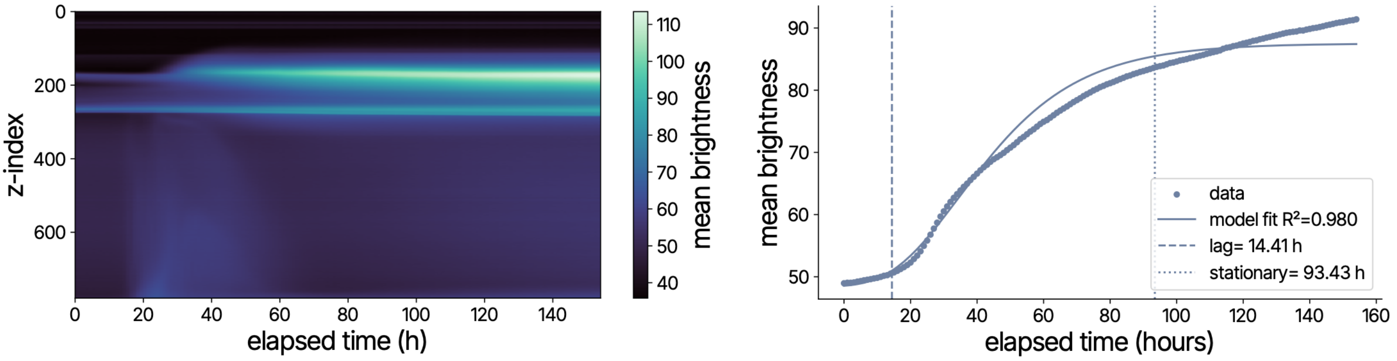


*Fig S48. Kymograph and growth curve of ROI 5 with membrane*

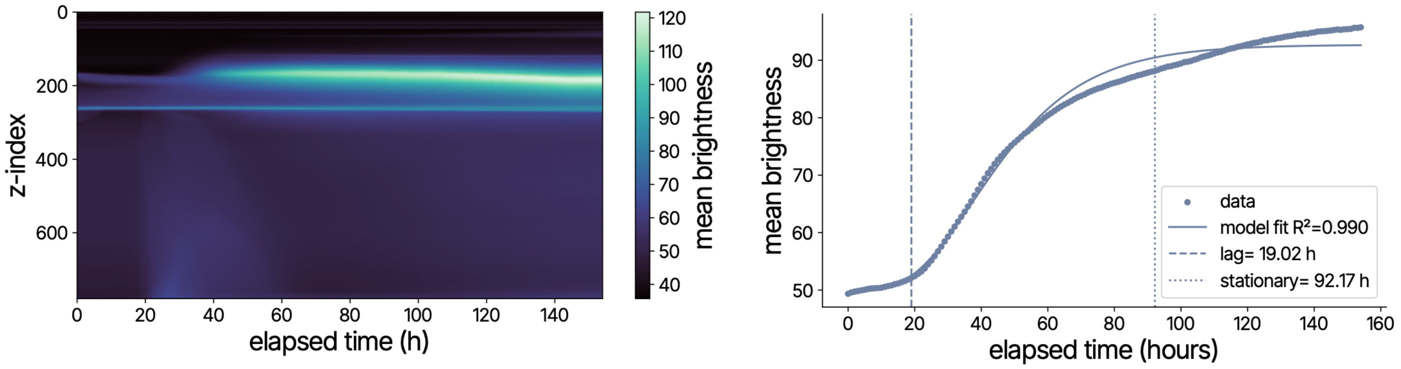


*Fig S49. Kymograph and growth curve of ROI 6 with membrane*

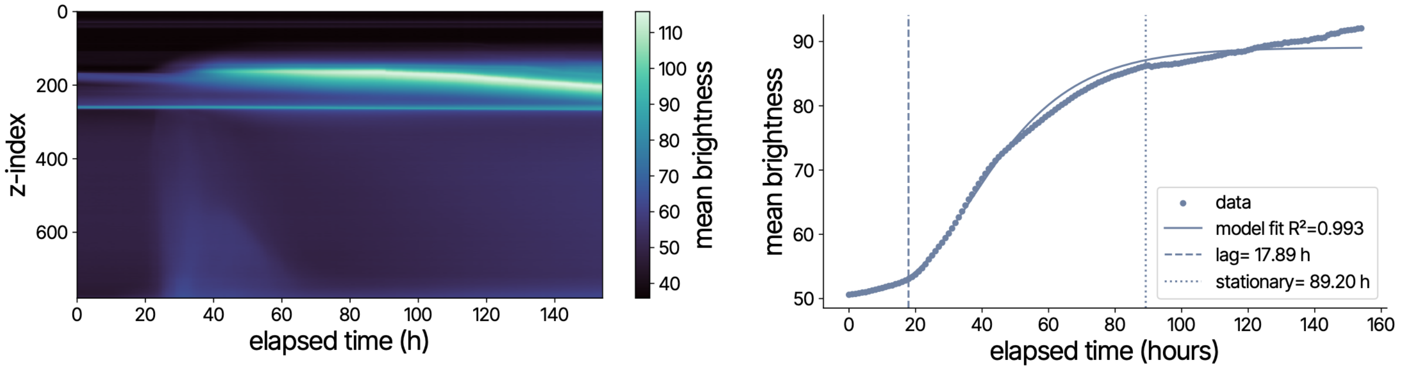


*Fig S50. Kymograph and growth curve of ROI 7 with membrane*

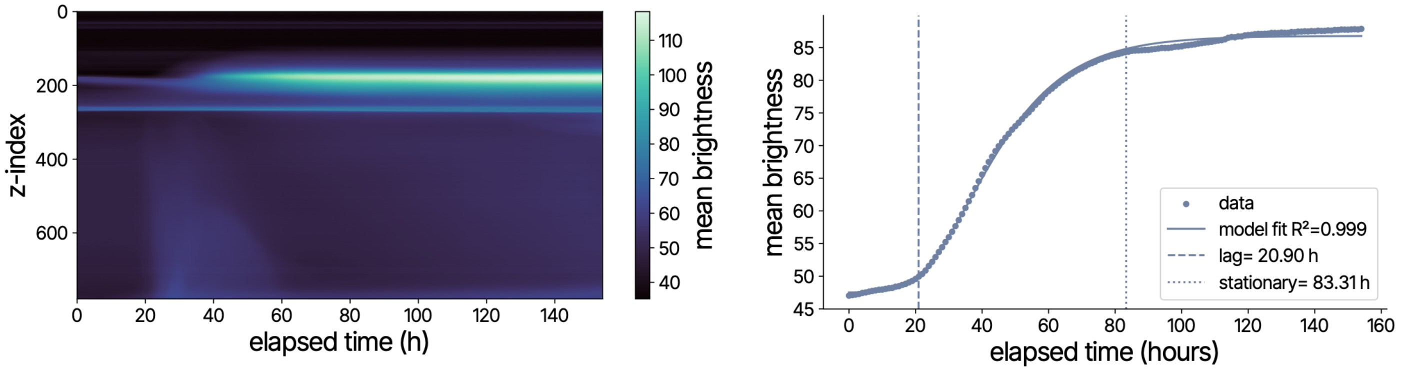


*Fig S51. Kymograph and growth curve of ROI 8 with membrane*

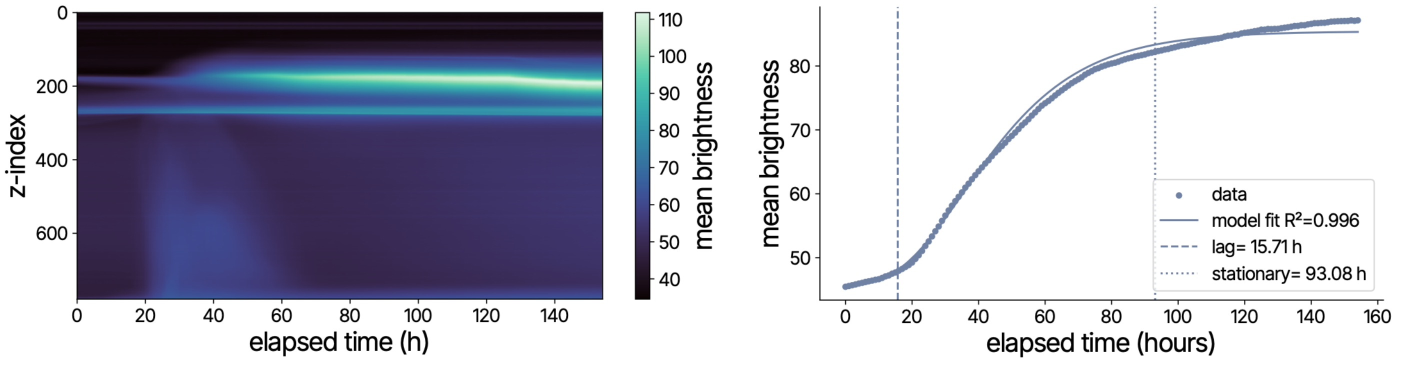


*Fig S52. Kymograph and growth curve of ROI 9 with membrane*

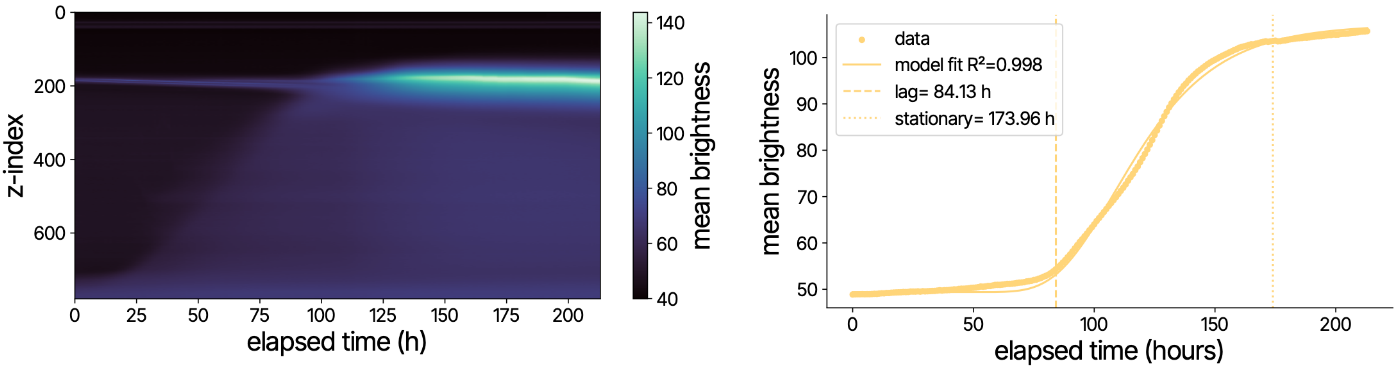


*Fig S53. Kymograph and growth curve of ROI 10 without membrane*

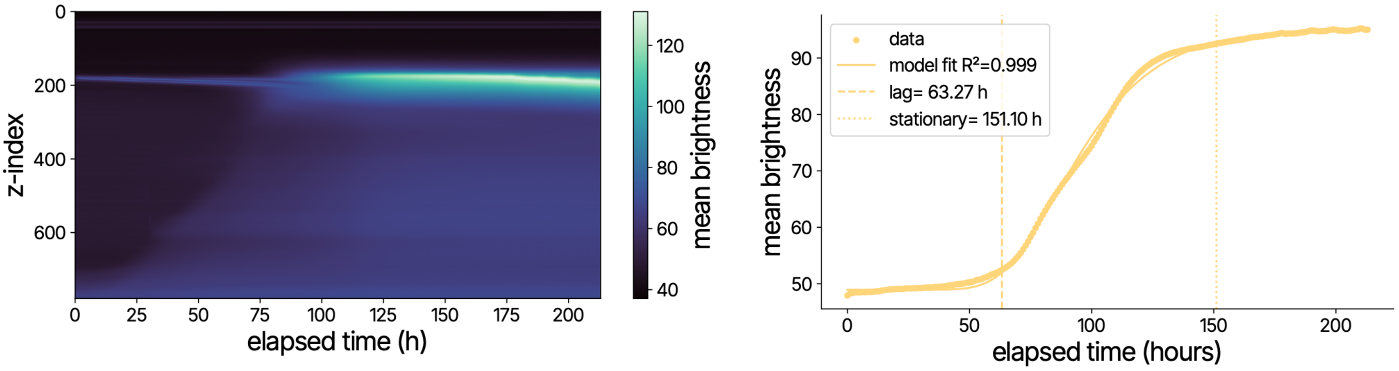


*Fig S54. Kymograph and growth curve of ROI 11 without membrane*

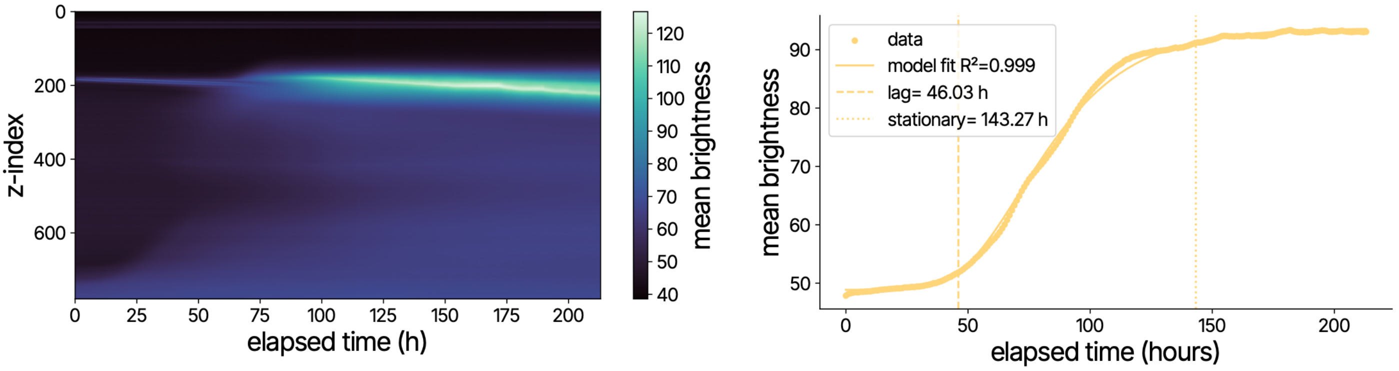


*Fig S55. Kymograph and growth curve of ROI 12 without membrane*

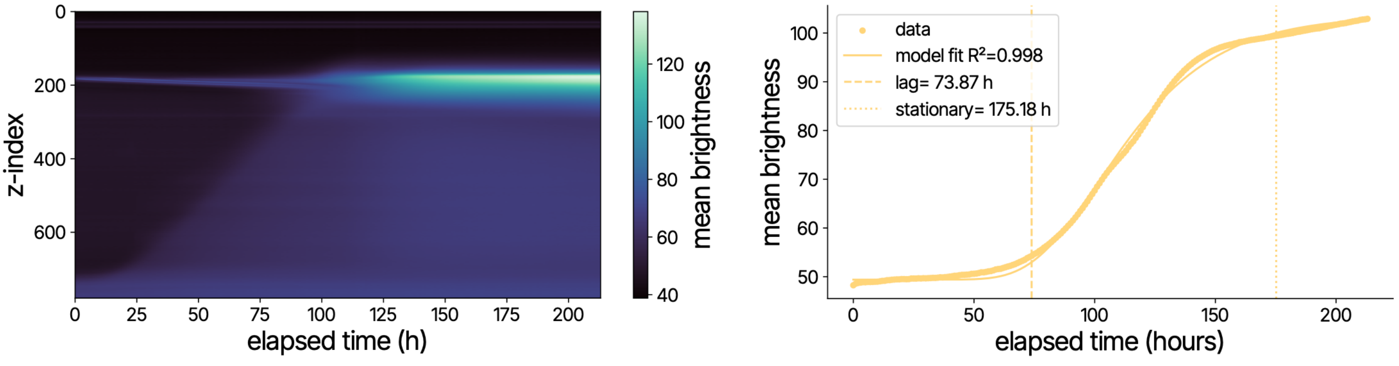


*Fig S56. Kymograph and growth curve of ROI 13 without membrane*

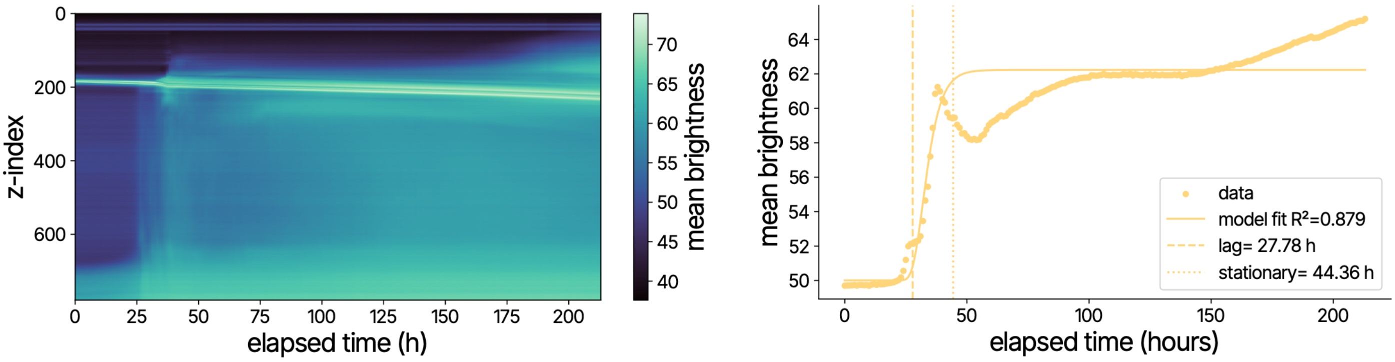


*Fig S57. Kymograph and growth curve of contaminated ROI 14 without membrane*

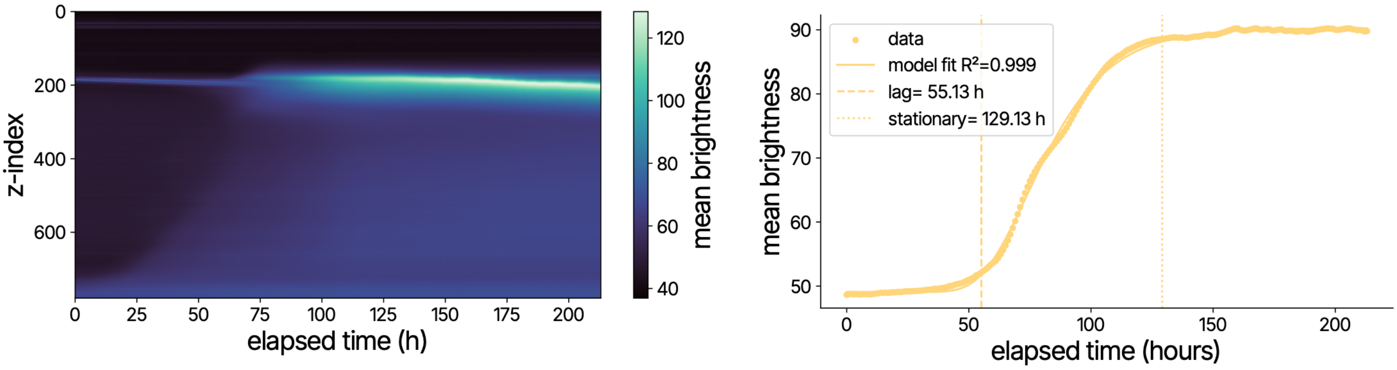


*Fig S58. Kymograph and growth curve of ROI 15 without membrane*

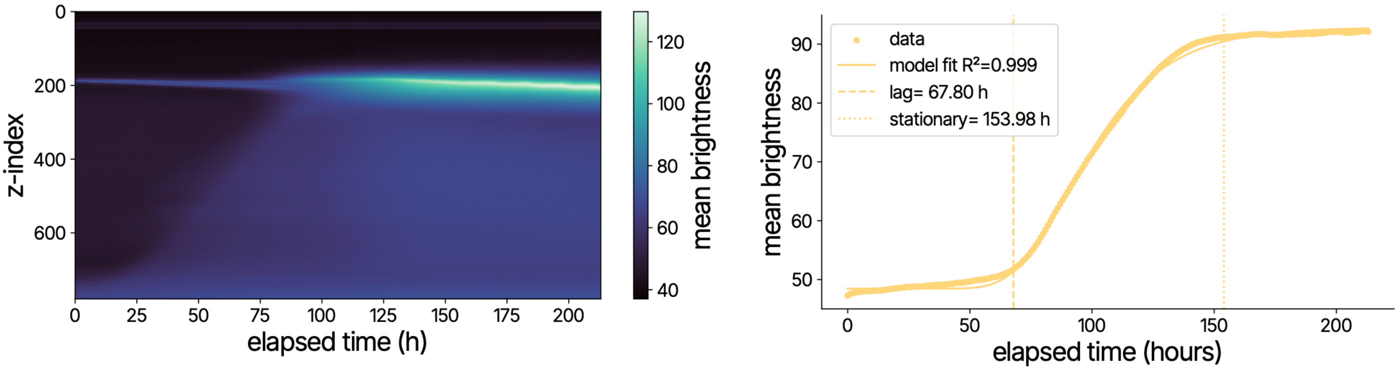


*Fig S59. Kymograph and growth curve of ROI 16 without membrane*

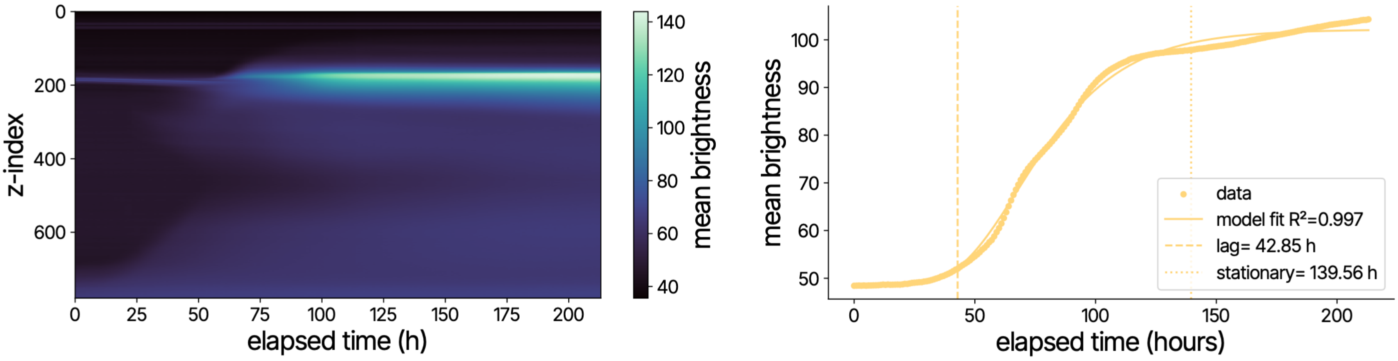


*Fig S60. Kymograph and growth curve of ROI 17 without membrane*

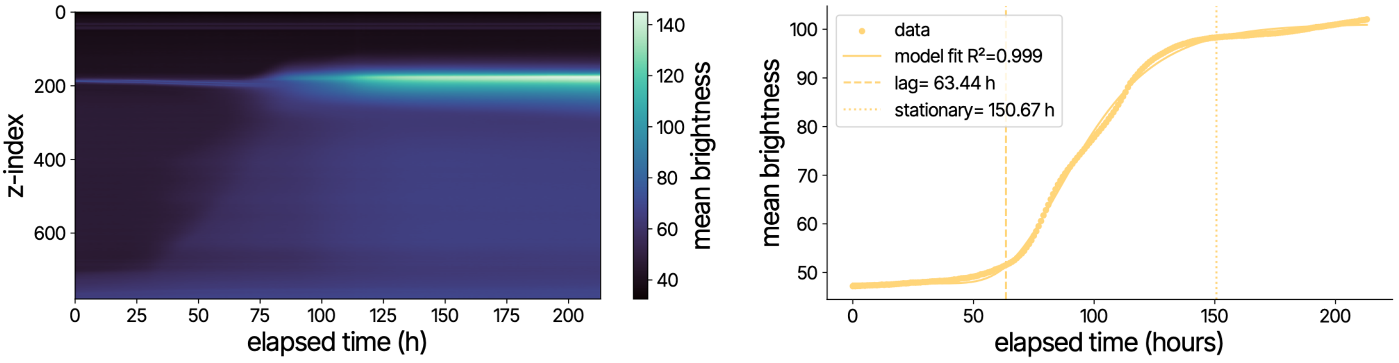


*Fig S61. Kymograph and growth curve of ROI 18 without membrane*

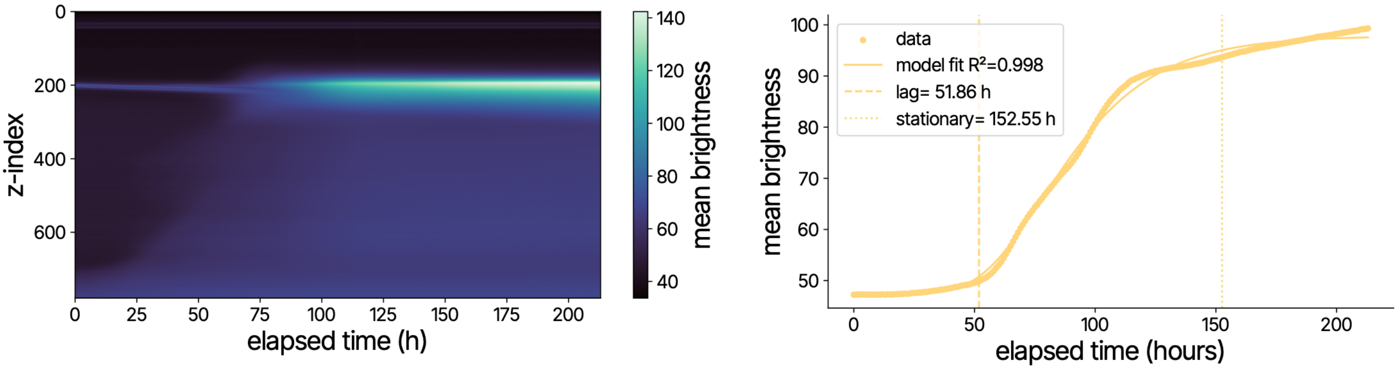


*Fig S62. Kymograph and growth curve of ROI 19 without membrane*

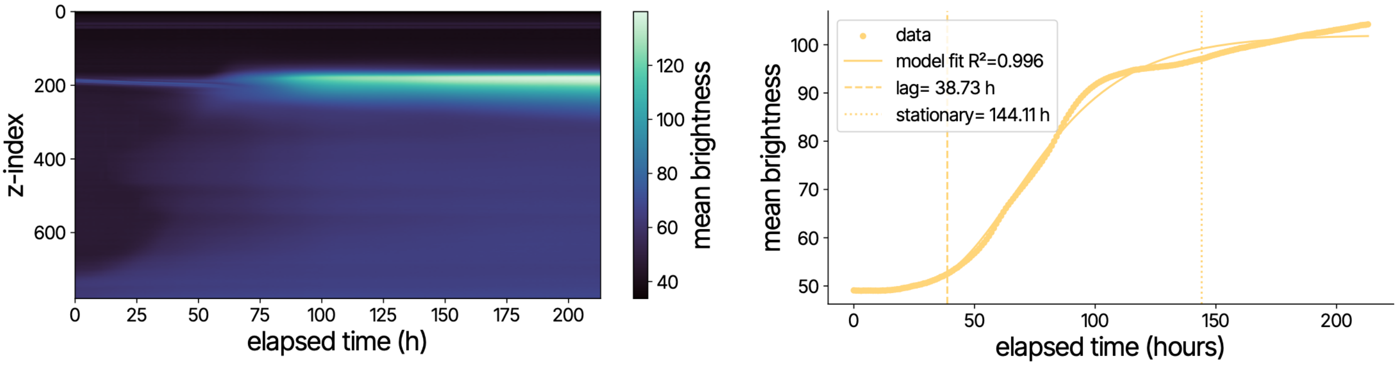


*Fig S63. Kymograph and growth curve of ROI 20 without membrane*
